# Progressive polysomal association of Upf1, Upf2, and Upf3 as ribosomes traverse mRNA coding regions

**DOI:** 10.1101/2021.12.03.471156

**Authors:** Robin Ganesan, Kotchaphorn Mangkalaphiban, Richard E. Baker, Feng He, Allan Jacobson

## Abstract

Upf1, Upf2, and Upf3 are the central regulators of nonsense-mediated mRNA decay (NMD), the eukaryotic mRNA quality control pathway generally triggered when a premature termination codon is recognized by the ribosome. The NMD-related functions of the Upf proteins likely commence while these factors are ribosome-associated, but little is known of the timing of their ribosome binding, their specificity for ribosomes translating NMD substrates, or the nature and role of any ribosome:Upf complexes. Here, we have elucidated details of the ribosome-associated steps of NMD. By combining yeast genetics with selective ribosome profiling and co-sedimentation analyses of polysomes with wild-type and mutant Upf proteins, our approaches have identified distinct states of ribosome:Upf association. All three Upf factors manifest progressive polysome association as mRNA translation proceeds, but these events appear to be preceded by formation of a Upf1:80S complex as mRNAs initiate translation. This complex is likely executing an early mRNA surveillance function.

## INTRODUCTION

Nonsense-mediated mRNA decay (NMD) is a eukaryotic translation-dependent mRNA quality control pathway whose central regulators are the three Upf proteins, Upf1, Upf2, and Upf3^1, 2^. NMD is initiated in response to atypical translation termination events, e.g., when an elongating ribosome encounters a termination codon that occurs prematurely within an open reading frame or in a context that otherwise renders termination inefficient^3, 4^. Although NMD’s existence has been known for decades^5–8^, and the pathway has been studied extensively^1, 2^, the precise mechanism by which the Upf proteins recognize a ribosome undergoing an atypical termination event and respond to it by triggering accelerated degradation of the associated mRNA remains unknown.

Several observations have suggested that the NMD functions of Upf1, Upf2, and Upf3 are exercised while these factors are bound to ribosomes. These include experiments demonstrating: 1) co-sedimentation of the Upfs with polyribosomes^9–11^, 2) interactions of the Upfs with release factors and ribosomal proteins^11–14^, and 3) roles for Upf1 and Upf3B in post-termination ribosome reutilization^15–17^. In contrast, several other studies have indicated that the central NMD factor Upf1 binds mRNA, but is displaced from coding regions by elongating ribosomes, and that the displaced factor principally resides on mRNA 3’-UTRs while translation is ongoing^18–20^. The latter observations imply that Upf1 responds *in trans* to an NMD-activating event, possibly as part of a “two-factor authentication” process for NMD that minimizes the likelihood that NMD activation will occur inadvertently^21^.

Regardless of where the Upf factors reside while awaiting their activation it appears that the initial steps in which they are committed to NMD functions are likely to be localized to elongating ribosomes. Hence, we have undertaken two approaches to examine the timing and roles of Upf1, Upf2, and Upf3 association with 80S ribosomes. In the first, we repeated older approaches in which the association of Upfs with ribosomes is examined on polysome gradients, but we have enhanced the sensitivity of such studies by including analyses of multiple mutants defective in different steps of NMD. In the second, we employed selective profiling of ribosomes respectively bound by Upf1, Upf2, or Upf3, and carried out these analyses as well in strains harboring NMD-altering mutations. In both approaches we sought to parse the steps between Upf factor ribosome binding and commitment of the associated mRNA substrate to NMD. Our approaches have allowed us to elucidate the timing and specificity of Upf factor association with translating ribosomes and ribosome bound NMD substrates, identify distinct states of ribosome association by Upf1 and its binding partners, and define a ribosome:Upf1 complex that appears to comprise an early step in NMD.

## RESULTS

### Mutations in *UPF1* Inhibit NMD and Alter Upf1 Polysome Association *in vivo*

To initiate an analysis of the interplay of translation and NMD, we first investigated the NMD phenotypes and polysome association of yeast Upf1 protein harboring amino acid substitutions that altered the protein’s enzymatic or RNA-binding activities. Wild-type (WT) *UPF1* and several well-characterized *upf1* alleles (see Extended Data Figure 1 for a map of the alleles analyzed), tagged at their C-termini with the FLAG epitope, were expressed on centromeric or episomal yeast plasmids and introduced into *upf1Δ* cells. Northern blotting analysis of *CYH2* pre-mRNA stabilization reiterated earlier observations ^12, 22–24^ that centromeric *C62Y*, *K436E*, and *DE572AA* alleles were defective in NMD to an extent comparable to *upf1Δ* (Figures 1a and 1b, top panels). Centromeric *R779C* and *RR793AA* alleles displayed partial NMD phenotypes, whereas the *AKS484HPA* mutation did not alter *CYH2* pre-mRNA abundance relative to centromeric WT *UPF1*. Expression of the mutant alleles on episomal plasmids resulted in partial restoration of NMD in *C62Y*, *K436E*, and *DE572AA* strains, but yielded no change in the NMD phenotypes of cells expressing the *R779C* or *RR793AA* alleles (Figures 1a and 1b, bottom panels). Association of the respective Upf1 mutant proteins with translating ribosomes was analyzed by western blotting of cytoplasmic extracts that had been fractionated by centrifugation through sucrose gradients (Figures 1c and 1d). Compared to WT *UPF1*, the *C62Y*, *R779C*, and *RR793AA* alleles resulted in reduced Upf1 recovery on polysomes when expressed from a centromeric plasmid, whereas polysome association was retained when *K436E*, *AKS484HPA*, and *DE572AA* mutant proteins were expressed from the same plasmid (Figure 1d, top, Figure 1e). Episomal expression of the *C62Y* and *RR793AA* alleles increased the polysome association of the respective Upf1 proteins (Figure 1d, bottom). Collectively, the results of Figure 1 show that: 1) *upf1* mutations can inhibit NMD function and reduce polysomal association of Upf1, 2) for some *upf1* alleles both activities can be compensated to some extent by Upf1 overexpression (e.g., *C62Y*), whereas for others (e.g., *RR793AA*) overexpression can improve Upf1 polysomal association without any effect on NMD activity, and 3) abundant polysome association of Upf1 is not necessarily correlated with NMD competence (e.g., *K436E* and *DE572AA*). These observations suggest that ribosome association of Upf1 is required but is not itself a triggering event in NMD.

**Figure 1.**
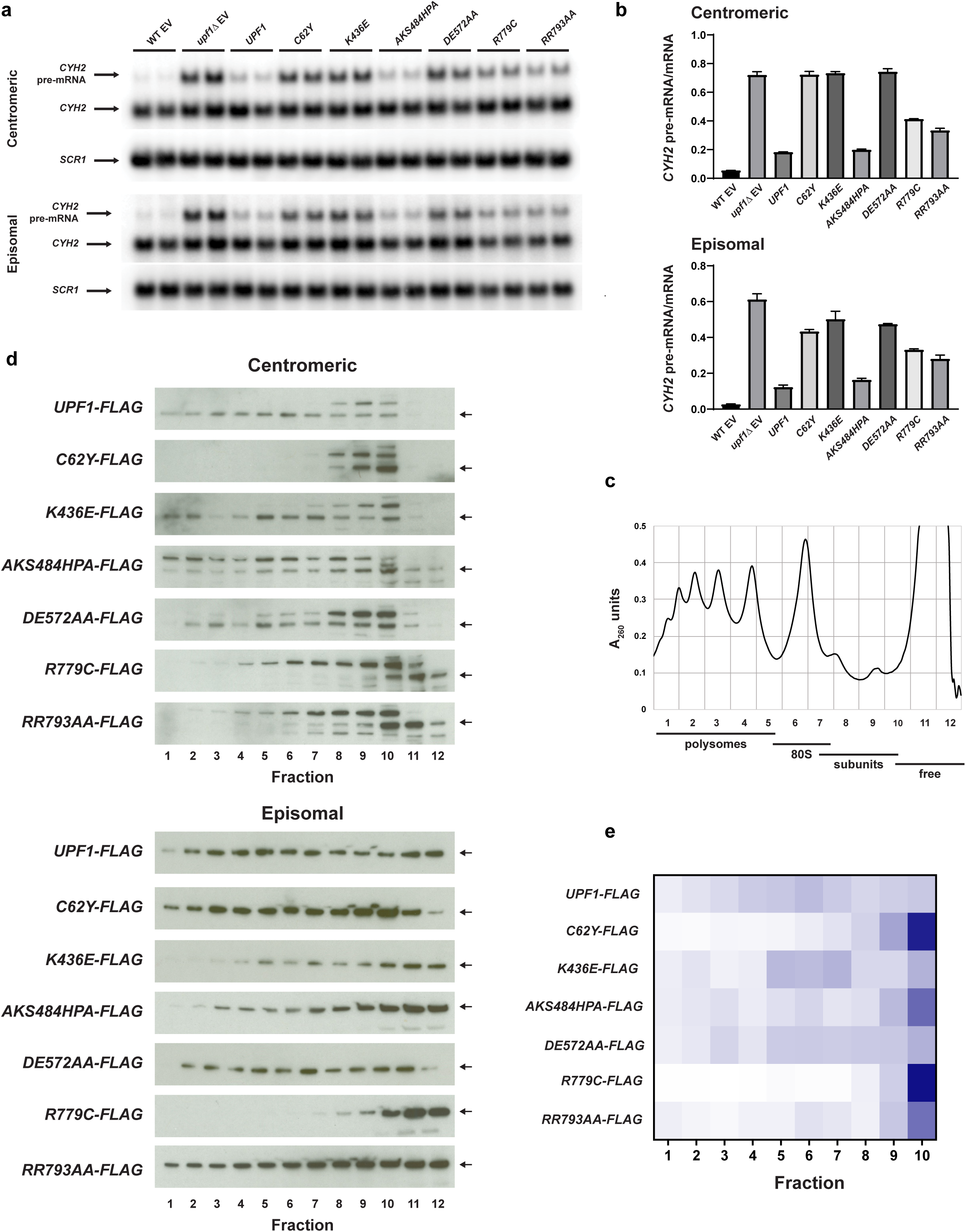
NMD phenotypes and polysome association of *UPF1* mutants. **a,** Northern blots of *CYH2* mRNA and pre-mRNA isolated from WT or *upf1Δ* cells containing an empty vector (EV) or from *upf1Δ* cells expressing *UPF1-FLAG* or its alleles from centromeric (top panel) or episomal vectors (bottom panel). Blots were probed for *SCR1* as a loading control. **b,** Ratio of *CYH2* pre-mRNA:mRNA for the strains depicted in part **a**. WT EV, wild-type *UPF1* strain containing empty vector; *upf1Δ* EV, *upf1Δ* strain containing empty vector, mean and range. **c,** Representative polysome trace from a *upf1Δ* strain containing centromerically expressed Upf1-FLAG across a 7-47% (right to left) sucrose gradient. **d,** Polysomal distributions of wild-type and mutant forms of Upf1. Lysates were prepared from a *upf1Δ* strain containing centromeric (top panel) or episomal (bottom panel) yeast plasmids expressing alleles of *UPF1-FLAG*. Western blots were probed with α-FLAG antibody. Arrow denotes expected migration of Upf1 protein; other bands in the top panel are background. **e,** Heat map of the mean Upf1-FLAG signal as percent total per fraction across sucrose gradient fractions 1-10 for centromeric WT *UPF1* and *upf1* mutant alleles from 3 (*C62Y-FLAG*, *K436E-FLAG*, *DE572AA-FLAG*, *R779C-FLAG*, *RR793AA-FLAG*) or 4 (*UPF1-FLAG*, *AKS484HPA-FLAG*) replicates per strain.

### Distinct States of Polysome Association by Upf1 and its Binding Partners

Upf1 and Upf2 are interacting proteins^25^ and the polysomal association patterns of the respective FLAG-tagged proteins expressed from episomal vectors are consistent with their known interactions. As shown in Figure 2a, FLAG-tagged Upf1 or Upf2 co-sediment throughout most of the polysomal fractions, being largely absent from only the heaviest polysomes. This co-sedimentation pattern changes, however, when either of the genes encoding the respective proteins is deleted. In *upf2Δ* cells, the distribution of FLAG-tagged Upf1 becomes skewed to the heaviest polysomes and, in *upf1Δ* cells, the distribution of FLAG-tagged Upf2 becomes skewed to lighter fractions (Figure 2a). These results suggest distinct ribosome-associated functional states for Upf1 and Upf2 and further confirm that, in steady-state, Upf1 and Upf2 interact. Dcp2, the catalytic component of the decapping enzyme, is also found on translating ribosomes^26^, and the co-translational recruitment of Dcp2 to NMD substrates is likely to occur via two highly similar Upf1-binding sites on Dcp2^27^. To address whether ribosome-associated Upf1 plays a role in recruitment of Dcp2, centromeric *UPF1-FLAG* was expressed in strains containing genomic HA-tagged *DCP2* or the *HA-dcp2-U1D-U2D* allele in which the two Upf1 binding sites have been deleted^28^. Sucrose gradient fractionation and western blotting demonstrated that deletion of the Upf1-binding sites on Dcp2 slightly shifted Dcp2 to lighter fractions (Figure 2b). This result suggests that, in steady-state, Upf1 recruits Dcp2 to mRNA on polysomes and enhances the overall sedimentation of Dcp2 as a Dcp2:Upf1:ribosome complex. Further, the combined results of Figures 1 and 2 validate the notion that Upf1 functions on polysomes and indicate that fully functional Upf1 can be associated with polysomes in at least four distinct states: bound or unbound to Upf2 and bound or unbound to Dcp2.

**Figure 2.**
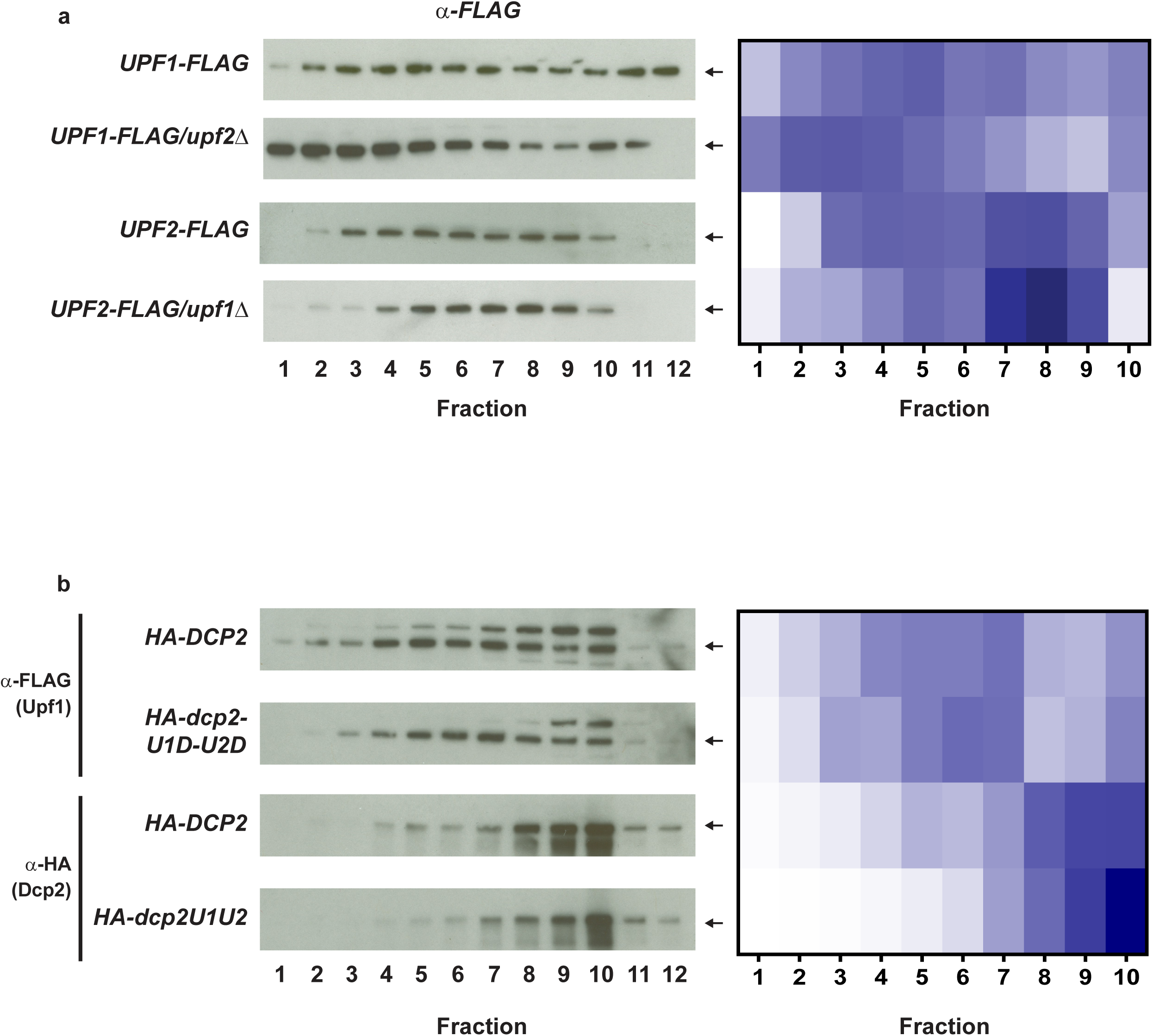
Polysomal distribution of Upf1, Upf2, and Dcp2 is modulated by their binding partners. **a,** Left, top two panels, polysomal distribution of episomally expressed Upf1-FLAG in *upf1Δ* or *upf1Δ*/*upf2Δ* strains; bottom two panels, distribution of episomally expressed Upf2-FLAG in *upf2Δ* or *upf1Δ* /*upf2Δ* strains. Western blots were probed with α-FLAG antibody. Arrow denotes expected migration of FLAG-tagged protein. Right, heat map of the mean Upf1-FLAG or Upf2-FLAG signal as percent total per fraction across sucrose gradient fractions 1-10, from 3 replicates per strain. **b,** Left, distribution of Upf1-FLAG (top two panels) or HA-Dcp2 or HA-dcp2-U1D-U2D protein (bottom two panels) across sucrose gradients of lysates prepared from genomic tagged *HA-DCP2/upf1Δ* or *HA-dcp2-U1D-U2D/upf1Δ* strains expressing centromeric *UPF1-FLAG*. Arrow denotes expected migration of tagged protein. Blots were probed with α-FLAG (top) or α-HA (bottom) monoclonal antibodies. Right, heat map of the mean Upf1-FLAG or HA-Dcp2 signals across sucrose gradient fractions 1-10 in *HA-DCP2/upf1Δ* or *HA-dcp2-U1D-U2D/upf1Δ* strains as in part **a**, from 2 replicates per strain. The episomal Upf1-FLAG in WT image and quantitation in Figure 2a is the same as in Figure 1d and 1e.

### Purification of FLAG-Upf1-, FLAG-Upf2-, and FLAG-Upf3-associated Ribosomes Yields Stoichiometric Recovery of Upf Factors with Ribosomal Proteins

To elucidate the roles of ribosome-bound Upf proteins in NMD, we carried out selective ribosome profiling analyses^29^ of 80S ribosomes associated with each of the three FLAG-tagged Upf factors. Pilot experiments in which FLAG-tagged Upf factors were expressed from centromeric plasmids did not yield sufficient immunoprecipitated ribosomes for construction of substantive ribosome profiling libraries. Hence, episomally expressed *UPF* genes were used for all experiments. To consider the possibility that location of the FLAG epitope tag might influence the results of the experiments, we examined *UPF1* constructs that were identical except for localization of the FLAG epitope to the N- or C-terminus. Likewise, as additional variables, we also examined cells in which all three *UPF* genes were simultaneously overexpressed, cells that were or were not treated with cycloheximide (CHX) prior to harvesting, and cells expressing mutant forms of the *UPF* genes. Upf factor-associated ribosomes were immunopurified as described previously^30^ and input (total) and immunopurified (IP) ribosomes were analyzed by western blotting. As shown in Extended Data Figure 2, immunopurification yielded ribosomes with strong signals for both ribosomal protein Tcm1 (Rpl3) and the respective FLAG-tagged Upf factor.

Confirmation of the specificity of recovery of Upf factor-associated ribosomes followed from mass spectrometry and intensity-based absolute quantification (iBAQ)^31, 32^ analyses of two biological replicates of each of the three N-terminally tagged *UPF* strains harvested with or without prior CHX treatment. The average normalized iBAQ for each of the total and immunopurified *FLAG-UPF* samples was calculated and compared to each other in scatterplots (Figure 3). In both the presence and absence of CHX, immunopurification resulted in substantial enrichment of the respective Upf protein such that its individual abundance was stoichiometric with the most abundant ribosomal proteins in the samples (blue dots in Figure 3a), a result strongly suggestive of a functional association of Upf proteins and ribosomes. Specificity of the FLAG immunopurification procedure was demonstrated by the lack of Upf1 enrichment in immunopurification of 6XHis-tagged *UPF1* (Figure 3b, top). With the exception of the respective FLAG-tagged *UPF* gene and the plasmid selective marker, all genes in these strains were present at their normal copy numbers. Thus, it was not surprising that immunopurification of FLAG-Upf1 or FLAG-Upf3 did not lead to substantial co-recovery of interacting Upf factors or components of the decapping complex. The co-recovery of interacting Upf factors in a *FLAG-UPF1* strain in which both *UPF2* and *UPF3* were also episomally expressed further demonstrated that the lack of co-recovery of interacting Upf factors in the *FLAG-UPF1* and *FLAG-UPF3* samples was due to low abundance of endogenous Upf2 and Upf3 and not the consequence of the immunopurification procedure (Figure 3b). Notably, however, immunopurification of Upf2 recovered endogenous Upf1 (Figure 3a), a result consistent with these two factors interacting directly and Upf1 being the most abundant of the three Upf proteins^33^.

**Figure 3.**
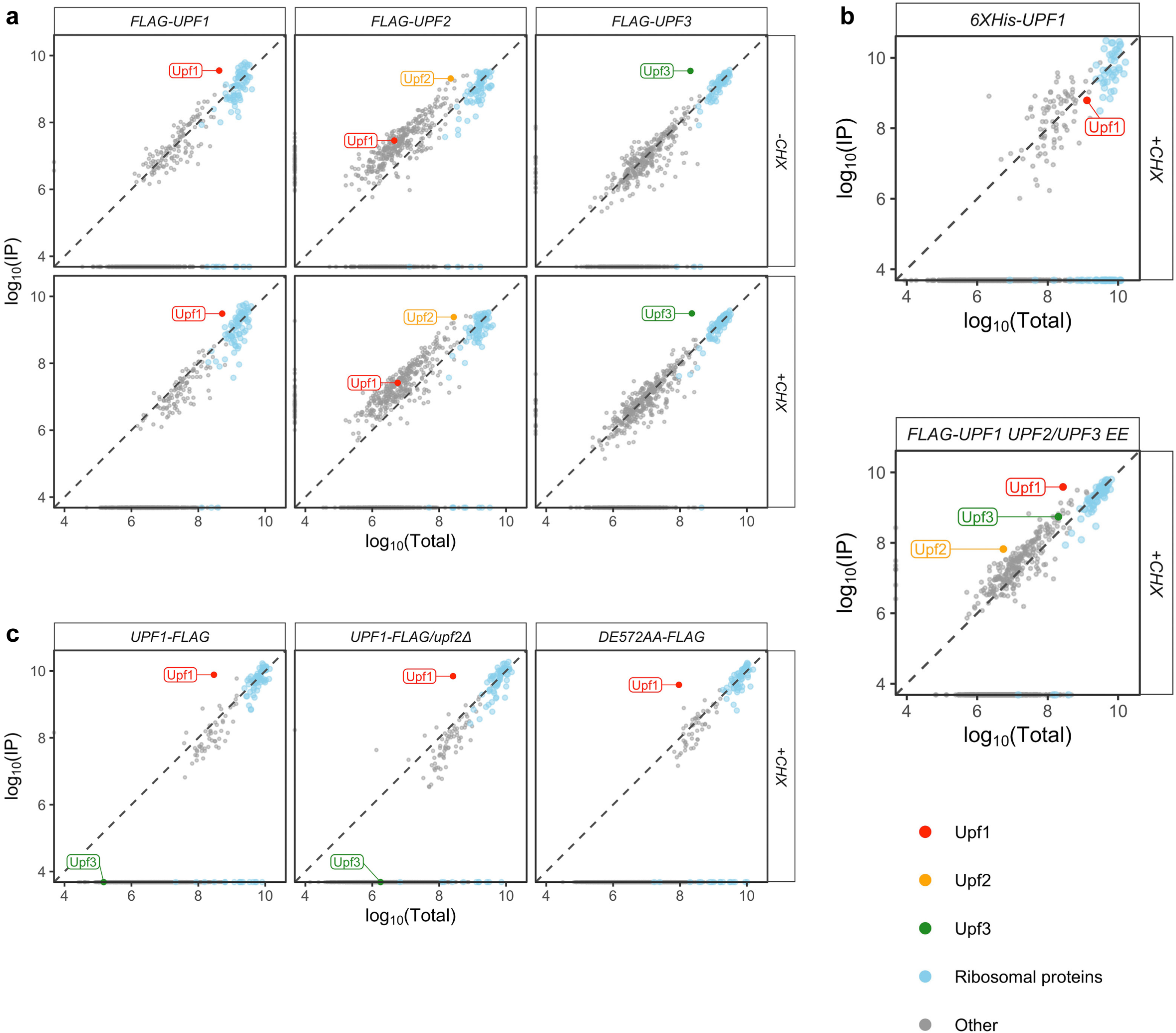
Stoichiometric recovery of FLAG-tagged Upf factors with ribosomal proteins. Input (Total) and immunopurified ribosomes (IP) from different lysates were analyzed by mass spectrometry. Intensity based absolute quantification (iBAQ) was performed and normalized across biosamples. Average iBAQ of biological replicates, log_10_-transformed, of proteins identified from IP were plotted against those from the Total ribosomes. Proteins with iBAQ of zero (cannot be log_10_-transformed) were plotted on top of the axes. Proteins above the diagonal dashed line are enriched in IP relative to Total, and proteins below the line are depleted in IP relative to Total. **a,** Ribosomes from lysates prepared with (+CHX) or without (-CHX) cycloheximide from cells expressing episomal *FLAG-UPF1*, *FLAG-UPF2,* or *FLAG-UPF3.* **b,** Ribosomes from cells expressing episomal *6XHis-UPF1* (top) or episomal *FLAG-UPF1* with *UPF2* and *UPF3* also episomally expressed (bottom). **c,** Ribosomes from cells expressing episomal *UPF1-FLAG* in a *UPF2* or *upf2Δ* background or *upf1*-*DE572AA-FLAG*.

Three strains expressing C-terminally tagged *UPF1* alleles (*UPF1-FLAG*, *DE572AA-FLAG* or *UPF1-FLAG*/*upf2Δ*) were treated with CHX and their ribosomes were also subjected to immunopurification. Mass spectrometry of the ribosomes immunopurified from three biological replicates of each of these strains also showed Upf1 enrichment that was stoichiometric with the ribosomal proteins in the samples (Figure 3c). A comparison of the extents of Upf1 recovery in the CHX-treated pellet samples of FLAG-Upf1 vs. Upf1-FLAG shows no significant difference in the respective extents of Upf1 recovery in the immunopurified samples, indicating that nascent Upf1, which would theoretically be enriched in the N-terminally tagged sample, makes only a modest contribution to overall Upf1 content of the samples. Importantly, inactivity of the NMD pathway or loss of Upf1 ATPase activity did not impair the purification of Upf1-associated ribosomes (compare the *UPF1-FLAG* sample to the *UPF1-FLAG/upf2Δ* and *DE572AA-FLAG* samples in Figure 3c).

### NMD Substrates are Enriched in Libraries from FLAG-Upf1-, FLAG-Upf2-, and FLAG-Upf3-associated Ribosomes

At some moment during translation of a nonsense-containing mRNA the association of Upf factors with a terminating ribosome should reflect commitment of that mRNA to NMD. Hence, we determined whether immunopurified Upf factor-associated ribosomes are enriched for ribosome footprints belonging to NMD substrates. DESeq2^34^ was employed to determine changes in ribosome footprint abundance from immunopurified Upf factor-associated ribosomes (IP) relative to input ribosomes (Total) for each mRNA. In all three *FLAG-UPF* strains processed without CHX treatment, the number of NMD substrates with increased recovery in IP relative to Total (log_2_(IP/Total) > 0) is proportionally higher than that of non-NMD substrates (Figure 4a, top, Extended Data Figure 3a, and Supplementary Table 1). NMD substrates are 1.4-1.6 times more likely to be found in the IP-enriched fraction than non-substrates and, among the IP-enriched fractions, are 1.9-2.6 times more likely to have > 2-fold enrichment than non-substrates (Supplementary Table 1, Fisher’s exact test). Logistic regression analysis on each sample also shows that a unit increase in log_2_(IP/Total) increases the odds of a transcript being an NMD substrate (Supplementary Table 1, Logistic regression). Additionally, the median log_2_(IP/Total) value of NMD substrates is significantly higher than that of non-substrates (Figure 4a, top, vertical dashed lines) and the latter are relatively depleted from all three IP samples. This enrichment of NMD substrates relative to non-substrate mRNAs confirms the trend seen when yeast Upf1 was immunopurified by a completely different procedure ^35^ and extends the NMD substrate enrichment observation to immunopurified Upf2- and Upf3-associated ribosomes. Although the degrees of NMD substrate enrichment relative to non-substrates are small (effect sizes < 0.2) (Figure 4a, top), it is expected because NMD substrates are rapidly degraded, i.e., the timeframe for detection is limited. Thus, consistent with the notion that stoichiometric recovery of Upf factors with a subset of ribosomes implies a functional relationship, we find that NMD substrates are enriched in all three sets of Upf factor-associated ribosomes. Repeating the same experiment with ribosomes isolated from cells subjected to brief CHX treatment yielded similar but slightly dampened results (Figure 4a, bottom, Extended Data Figure 3a). For this set of Upf factor associated ribosomes the magnitude of difference between NMD substrates and non-substrates, although significant, is reduced relative to the no-CHX samples evaluated in the same manner (compare effect sizes or odds ratio when all transcripts considered). The samples from CHX-treated cells also indicate that mutations in the NMD pathway affect NMD substrate recovery in the IP samples. Relative to the *UPF1-FLAG* IP samples, the *DE572AA* mutation enhances NMD substrate recovery whereas the *upf2Δ* mutation slightly diminishes NMD substrate recovery (Figures 4b and Extended Data Figure 3b, and Supplementary Table 1).

**Figure 4.**
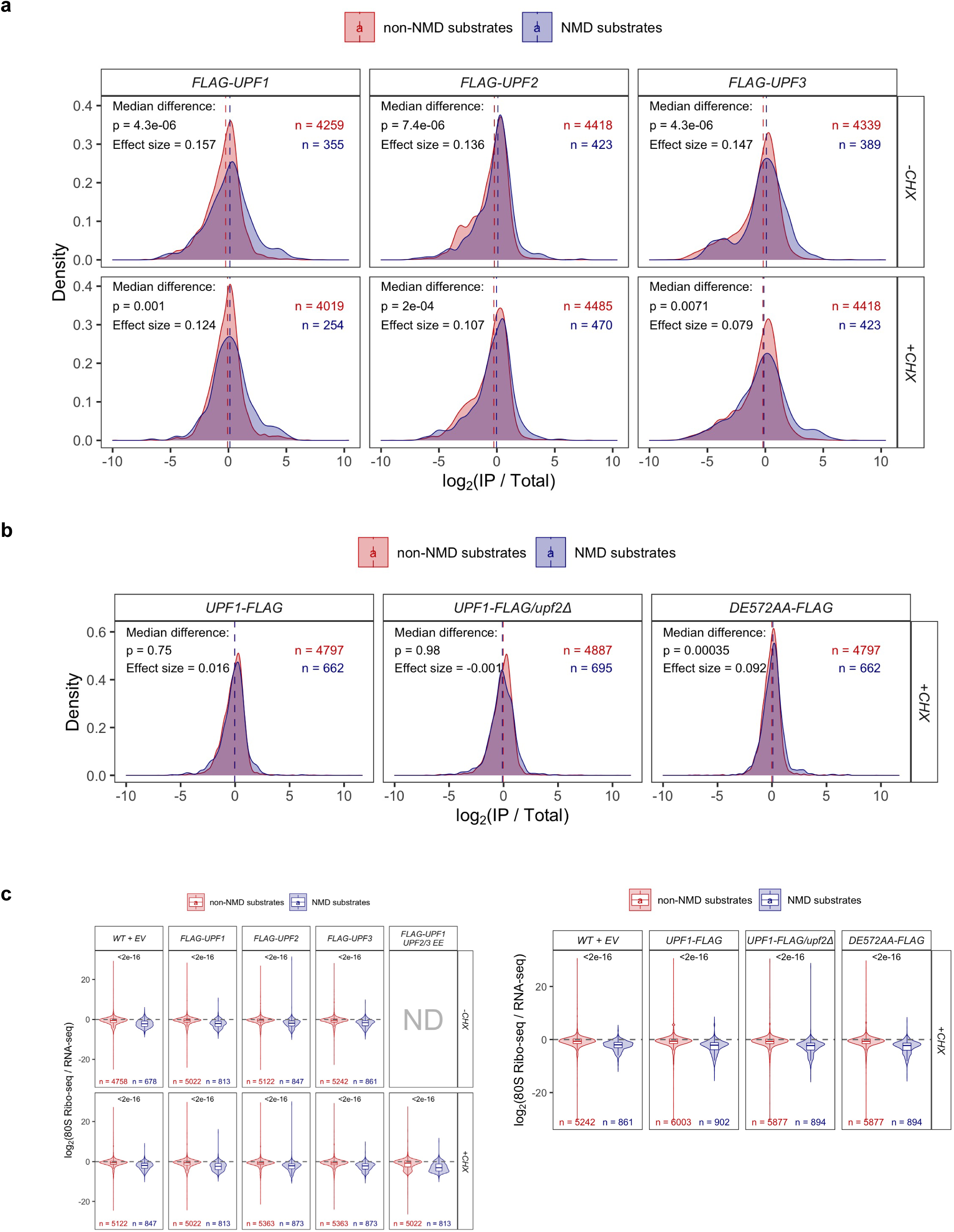
NMD substrates are enriched in Upf factor-associated ribosomes relative to total ribosomes. Relative translating ribosome footprint recovery in immunopurified (IP) vs. input (Total) ribosomes: log_2_(IP/Total) for **a,** *FLAG-UPF1*, *FLAG-UPF2,* or *FLAG-UPF3* and **b,** *UPF1-FLAG*, *UPF1-FLAG*/*upf2Δ*, or *upf1*-*DE572AA-FLAG* strains. Vertical dashed line indicates median log_2_(IP/Total). The effect size measures the degree of difference between median log_2_ fold change of NMD substrates and that of non-substrates. **c,** Translation efficiency, defined by abundance of translating ribosome footprints normalized to the corresponding mRNA abundance of NMD substrates (blue) and non-substrates (red). For all, significant differences between median log_2_ fold change of NMD substrates and that of non-substrates was determined by Wilcoxon’s rank sum test with Benjamini-Hochberg method for multiple testing correction.

Our ability to quantitate ribosome footprints from NMD substrates and non-substrates allowed us to revisit an earlier observation indicating that NMD substrates are poorly translated relative to non-NMD substrates^36^. We confirmed this observation in each of the 13 distinct ribosome profiling datasets generated in this study, finding that NMD substrates consistently have lower translational efficiency than non-NMD substrates even in those cases where NMD has been inactivated (Figure 4c).

### Upf1 Association with 80S Ribosomes in CHX-treated Cells Promotes the Formation of Atypical Ribosome-protected mRNA Fragments

To gain insight into the relationship between Upf factors and translating ribosomes, we examined the nature of the ribosome protected fragments recovered in *FLAG-UPF* libraries. Analyses of ribosome protected footprint length distribution in libraries from cells without CHX treatment showed that footprints of ∼20-23 nt in length (“small” size, hereafter denoted as “S”) were predominant, followed by ∼27-32 nt footprints (“medium” size, hereafter denoted as “M”) (Figure 5a, green lines). These previously detected^37^ footprint sizes were present in cells with or without FLAG-tagged *UPF* genes and in samples with or without prior immunopurification of Upf factor-associated ribosomes. In the presence of CHX, the predominant footprint sizes from total or immunopurified ribosomes were the M size class; this is expected since CHX is known to freeze ribosomes with occupied A sites, yielding footprints ∼28 nt in length ^37, 38^ (Figure 5a, red lines). Notably, ribosomes from CHX-treated strains expressing *FLAG-UPF1*, but not those expressing *FLAG-UPF2* or *FLAG-UPF3*, yielded an atypical footprint size class of ∼37-43 nt (“large” size, hereafter denoted as “L”) (Figure 5a, red solid lines). The ratio of L:M footprints was increased in the *FLAG-UPF1* IP libraries compared to total libraries (Figure 5a, red solid lines), and the L footprints were not detectable in libraries prepared from cells without CHX treatment (Figure 5a, green lines).

**Figure 5.**
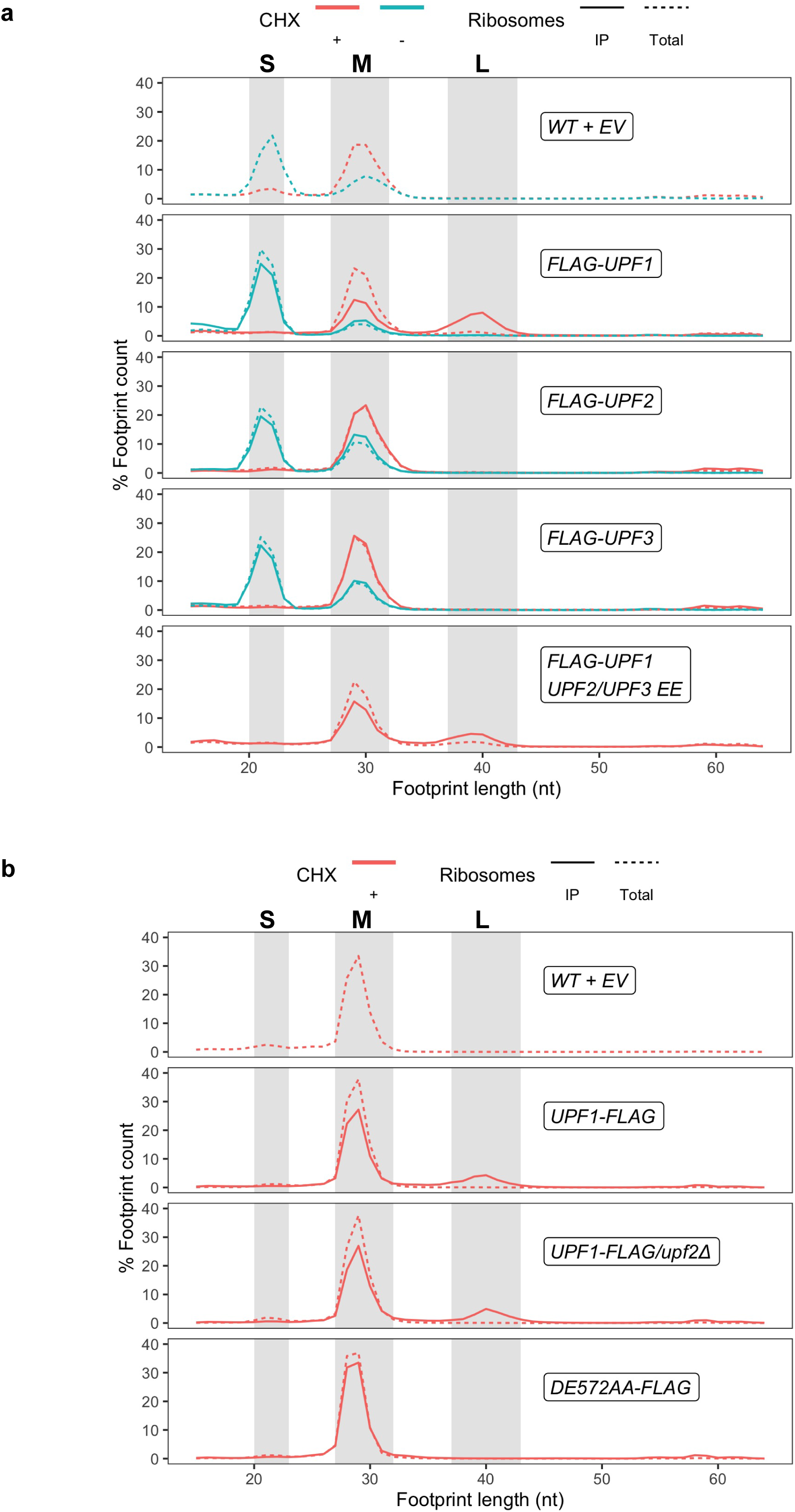
Distribution of footprint length for each strain and cycloheximide treatment condition. **a,** Total and IP ribosome profiling libraries from WT+EV, *FLAG-UPF1*, *FLAG-UPF2,* or *FLAG-UPF3* strains. **b,** Total and IP ribosome profiling libraries from WT+EV, *UPF1-FLAG*, *UPF1-FLAG*/*upf2Δ*, or *upf1*-*DE572AA-FLAG* strains. Fractions of each footprint length (nt) were calculated and averaged among replicates. Green and red lines indicate libraries prepared in the absence and presence of cycloheximide, respectively. Dashed and solid lines represent total and immunopurified (IP) ribosomes, respectively. Gray shaded areas highlight different classes of footprint size: from left to right, 20-23 nt = small (S), 27-32 nt = medium (M), and 37-43 nt = large (L).

As different classes of ribosome footprint sizes reflect distinct states of the translation elongation cycle^37^, we wondered whether different footprint classes might also represent distinct steps of NMD. If one size class represents a state of pre-commitment to NMD while another class represents a commitment to NMD, NMD substrates may show an increase in footprints of the latter relative to the former, while non-substrates may not. Accordingly, we calculated the ratio of the relative recovery of the minor footprint class over that of the major class for individual mRNAs. In all strains untreated with CHX, the changes in M over S footprint abundance between NMD substrates and non-substrates are not statistically different (Extended Data Figure 4, top panel). A similar conclusion is reached for the changes in L over M footprint abundance between NMD substrates and non-substrates in the *FLAG-UPF1* strain treated with CHX (Extended Data Figure 4, bottom panel). In all cases, most of the transcripts and the median log_2_(minor/major) are centered at 0, indicating that the proportion of major and minor footprints of most transcripts does not differ. Thus, different footprint size classes captured here are no more NMD-specific than one another.

We also considered the possibility that the atypical L footprints might be a consequence of non-stoichiometric expression of *UPF2* or *UPF3* relative to *UPF1* in cells expressing episomal *FLAG-UPF1,* but concurrent episomal expression of *UPF2, UPF3,* and *FLAG-UPF1* still yielded L footprints from immunopurified ribosomes (Figure 5a, bottom panel). Likewise, we considered the possibility that the L footprints were caused by the 5’-FLAG epitope on Upf1, but analysis of libraries generated from cells expressing *UPF1-FLAG* again showed recovery of the L footprints in immunopurified ribosomes (Figure 5b). Therefore, this unique class of footprints is most likely specific to Upf1 association with the 80S ribosome.

### Upf1 Function but not Full Activity of the NMD Pathway is Required for Formation of Atypical Footprints by Upf1-associated Ribosomes

To understand the origin of the L footprints, we determined whether their formation required Upf1 function, as well as function of the entire NMD pathway. Total and immunopurified ribosomes from CHX-treated *upf1Δ* strains expressing the *DE572AA-FLAG* allele, or *UPF1-FLAG* in *UPF2* or *upf2Δ* cells, were subjected to selective ribosome profiling and analysis of footprint length distribution (Figure 5b). The L footprints were undetectable in any libraries from total ribosomes but were recovered in libraries prepared from immunopurified ribosomes isolated from *UPF2* or *upf2Δ* cells expressing *UPF1-FLAG.* The latter result indicates that full functionality of the NMD pathway is not required to form L footprints and the former result confirms that a 5’ FLAG tag does not promote L footprint formation. However, the L footprints were undetectable in libraries prepared from the *DE572AA-FLAG* strain (Figure 5b), i.e., full Upf1 function is required for their formation. Upf1-*DE572AA* is present on polysomes (Figures 1d and 1e) and is able to interact with Rps26 in a two-hybrid assay ^12^; therefore, the loss of the L footprints in cells expressing *DE572AA-FLAG* is likely due to the inability of this mutant Upf1 to promote a ribosome-associated function that requires Upf1’s ATP hydrolysis activity.

### L Footprints are Generated by Additional Protection of mRNA 5’ to the Normal Ribosome-protected Segment

The recovery of L footprints from CHX-treated cells expressing *FLAG-UPF1* or *UPF1-FLAG* (Figures 5a and 5b) suggests two likely possibilities: either a fraction of Upf1 is bound to ribosomes in a configuration that inhibits RNase1 digestion of mRNA that usually occurs at the edge of the ribosome during library preparation or CHX treatment has induced a ribosome collision that results in Upf1 association with the resulting disome and NGD (No Go Decay)-like endonucleolytic cleavage in the A site of the colliding ribosome^39^. Mapping of the 5’ and 3’ ends of the M and L footprints recovered from Upf1-associated ribosomes using the start and stop codons as reference points showed that the 3’ ends but not the 5’ ends of M and L footprints are aligned at the same nucleotide location (Figure 6). Thus, the size difference between M and L footprints is entirely attributable to an extension on the 5’ side of the normal ribosome-protected fragment. The 5’ ends of the typical M footprints are ∼12-13 nt upstream from the reference codon, while the 5’ ends of the atypical L footprints are ∼23-25 nt upstream from the stop codon, regardless of the N- or C-terminal placement of the FLAG epitope, confirming that the extra 10-13 nt that extend “ribosome protection” do so on the 5’ side of the fragment.

**Figure 6.**
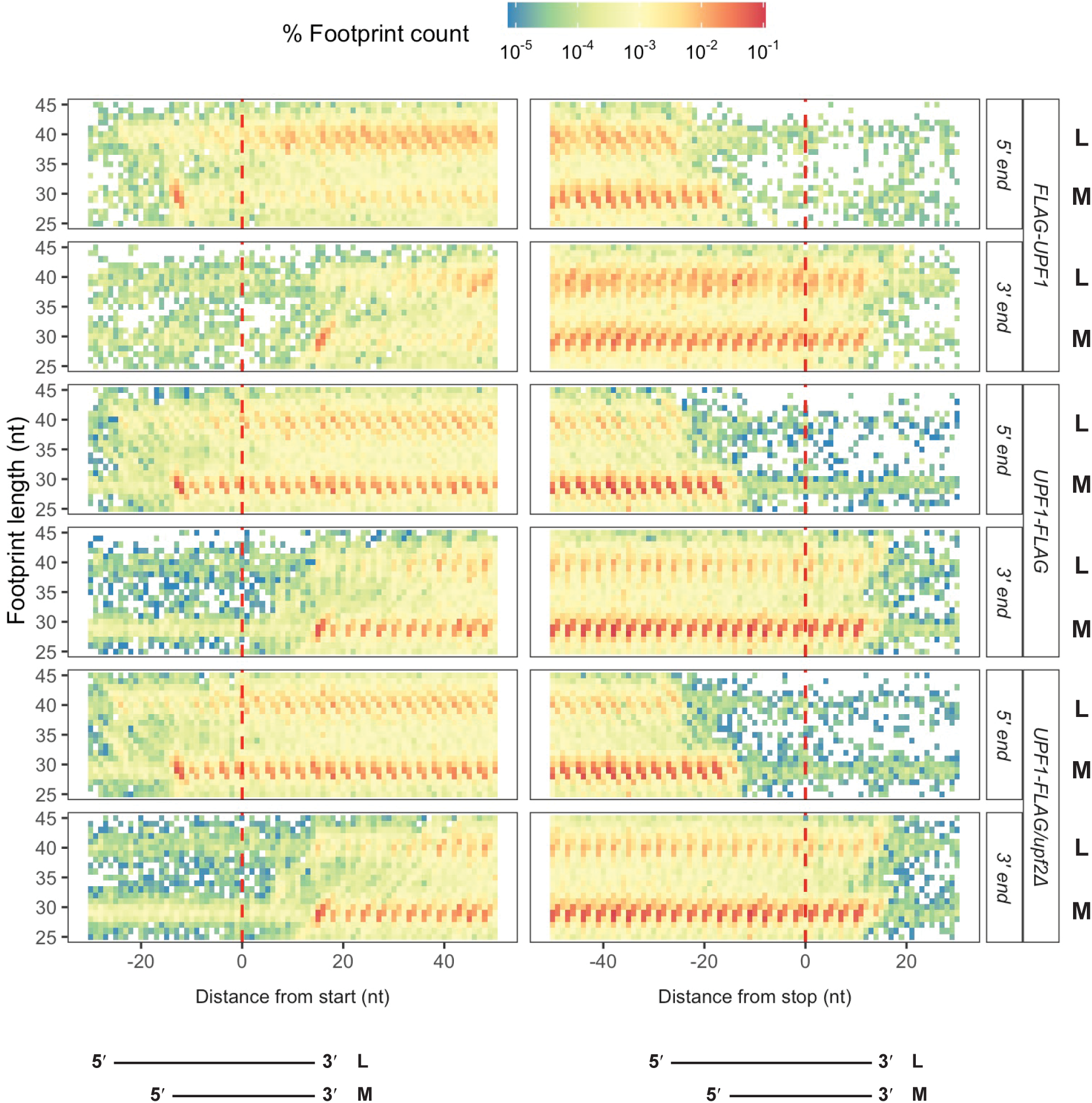
Atypically large footprints are attributable to a 5’ extension of the mRNA region protected by the ribosome. Mapping of the 5’ and 3’ ends of footprint lengths 25-45 nt relative to the start and stop codons from *FLAG-UPF1*, *UPF1-FLAG*, and *UPF1-FLAG/upf2Δ* (+CHX). Different footprint lengths align at the 3’ ends but diverge at the 5’ ends. L and M fragment size classes are as indicated for each sample. Lines at the bottom of the figure indicate the approximate 5’ and 3’ limits of the L and M fragments over the start and stop reference codons.

### L Footprints Reflect an Early Phase of Upf1 Association with the Ribosome

Metagene analyses comparing the distributions of S, M, and L ribosome-protected fragments across normalized coding regions from total and immunopurified ribosomes of *FLAG-UPF* strains show that, in all three strains analyzed in the absence of CHX, the S and M footprints from immunopurified ribosomes are markedly underrepresented in approximately the first half of the coding region and become overrepresented in the second half of the coding region compared to total ribosomes (Figure 7a, Extended Data Figure 5). The same pattern holds for the M footprints in all three *FLAG-UPF* libraries from CHX-treated cells (Figure 7b, top panel, Extended Data Figure 5). The metagene distribution of S and M footprints recovered from immunopurified ribosomes was nearly identical regardless of the presence or absence of CHX (Figure 7d, bottom panel), except for a small peak over the start codon in M footprints. This suggests that Upf1 is not stripped off the coding region by translating ribosomes^18–20^ and that Upf1 association with ribosomes occurs routinely during the course of translation elongation.

**Figure 7.**
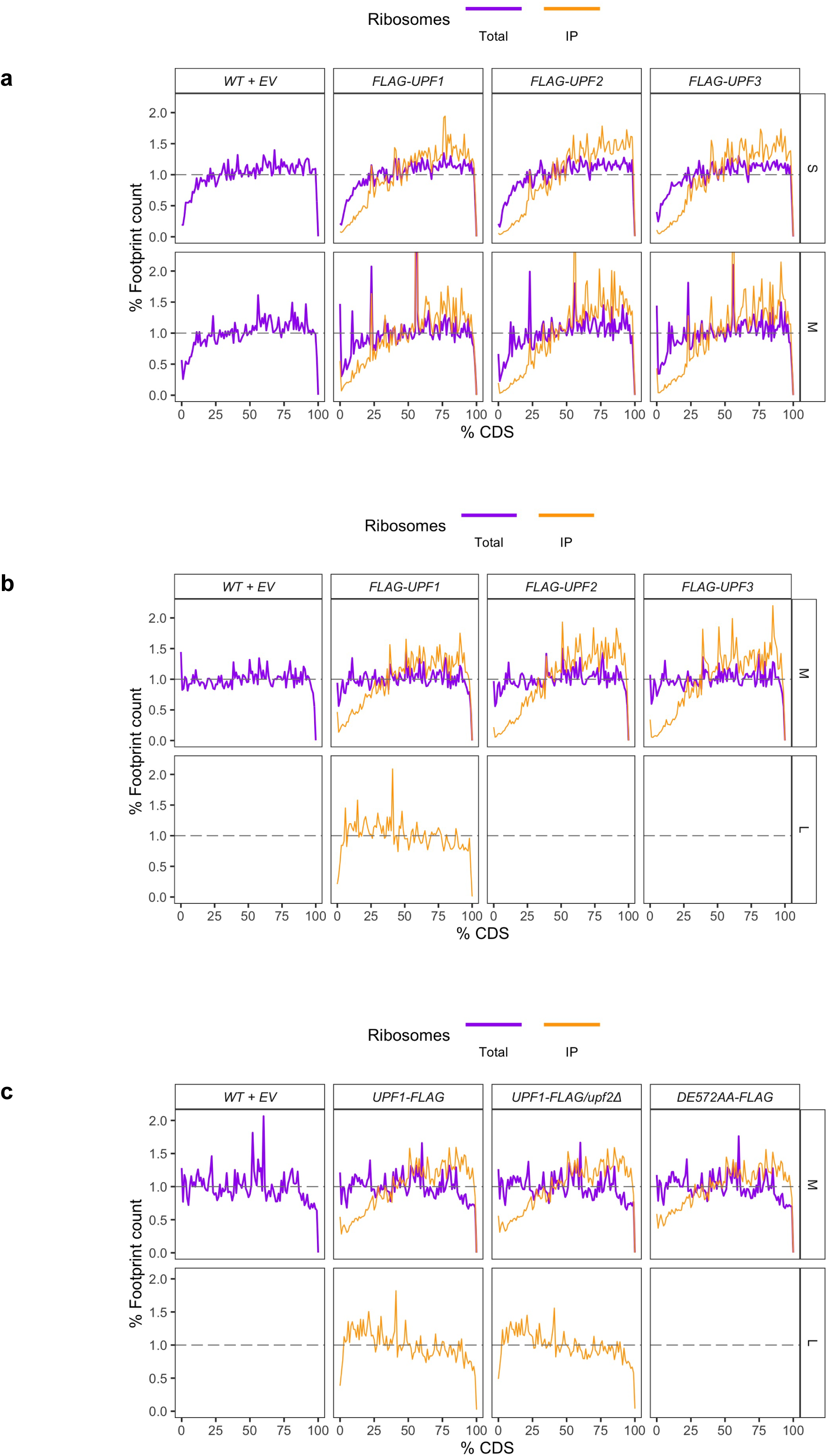
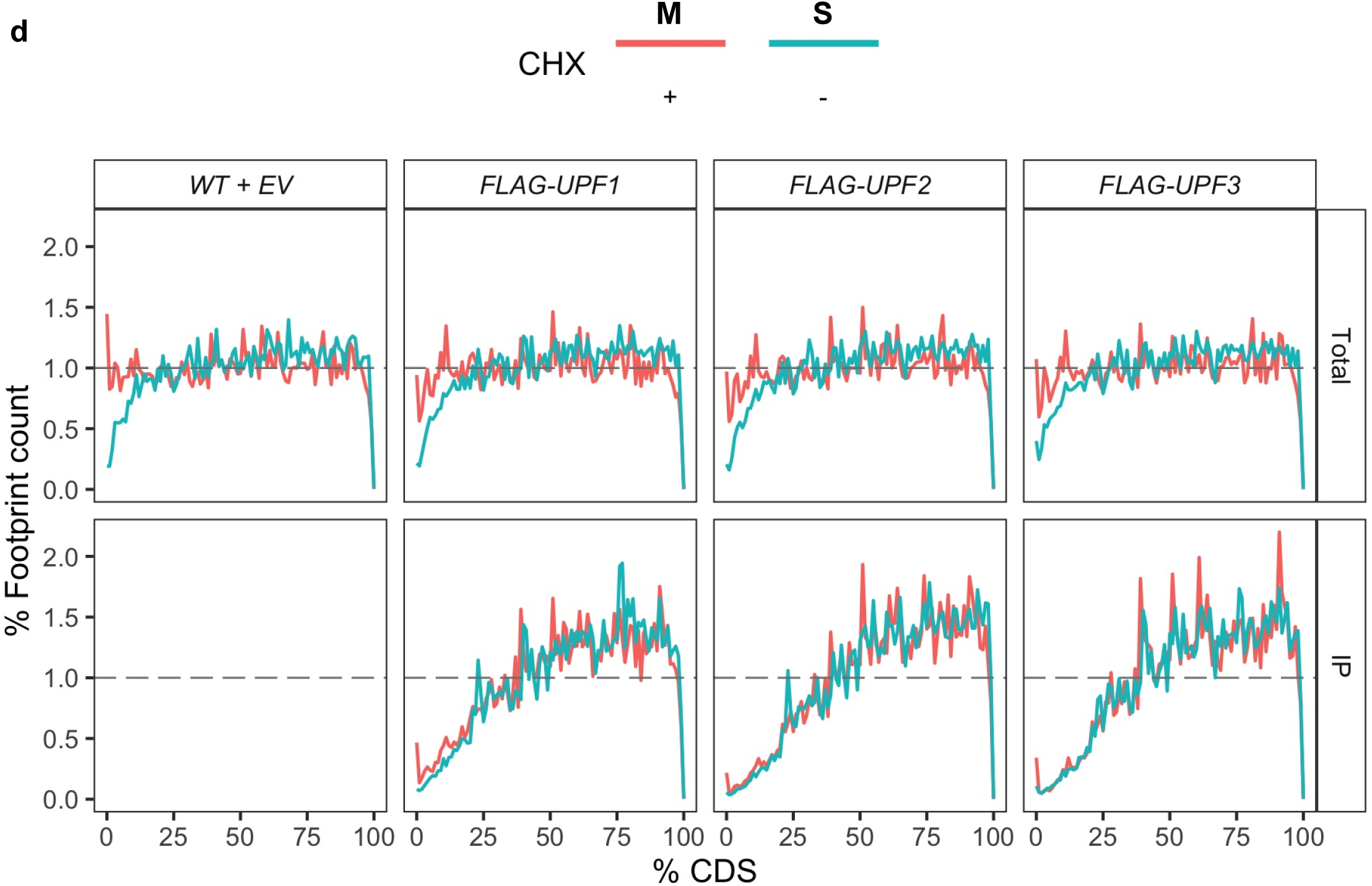
Upf factors progressively associate with ribosomes across mRNA coding regions except when Upf1 association forms L footprints. Distribution of footprint abundance across the coding region (CDS) for each footprint size class from total (purple) or IP (orange) ribosomes profiling libraries. **a,** Small and medium footprint distribution from strains expressing WT+EV, *FLAG-UPF1*, *FLAG-UPF2,* or *FLAG-UPF3*, all in the absence of CHX treatment. **b,** Medium and large footprint distribution from CHX-treated cells expressing WT+EV, *FLAG-UPF1*, *FLAG-UPF2,* or *FLAG-UPF3*. **c,** Medium and large footprint distribution from CHX-treated cells expressing WT+EV, *UPF1-FLAG*, *UPF1-FLAG*/*upf2Δ*, or *upf1-DE572AA-FLAG.* Each gene’s CDS was divided into 100 bins and the percentage of footprints’ P-sites belonging to each bin in all genes was calculated. Grey dashed line signifies a theoretical number where each percentage of CDS contains an equivalent number of footprints for a sum of 100% across all 100 bins. **d,** Distribution of dominant footprint (red = Medium, M for +CHX; green = Small, S for -CHX) abundance across the coding region. Data is the same as used in **a** and **b** top panels, but re-plotted in such a way to directly observe the differences or similarity in footprint distribution between the absence and presence of CHX during the library preparation of each strain.

The L footprints, which are only detectable with Upf1-associated ribosomes in CHX-treated cells, accumulate rapidly at the beginning of mRNA coding regions and slowly decrease their relative accumulation across the entire coding region in libraries prepared from immunopurified ribosomes from *FLAG-UPF1, UPF1-FLAG,* or *UPF1-FLAG/upf2Δ* strains (Figures 7b and 7c, Extended Data Figure 5). The accumulation of L footprints toward the 5’ end of mRNAs is significantly higher than that of M footprints (Supplementary Table 2). These patterns, and their expected 3-nt periodicity, are also evident in higher resolution metagene plots of the first and last 100 nt of the coding regions (Extended Data Figure 6). Notably, L footprints do not accumulate over the start codon as do M footprints (Extended Data Figures 5 and 6, green lines). The rapid appearance of L footprints relative to ribosome progression across mRNA coding regions (Figures 7, Extended Data Figure 6), as well as their comparable enrichment for NMD substrates to the typical ribosome footprints (Extended Data Figure 4), indicates that their formation is likely to be an early event in NMD when Upf1 is initiating its association with the ribosome.

## DISCUSSION

### Upfs Function While Associated with Ribosomes

Given the uncertainties underlying the steps between the association of Upf factors with elongating ribosomes and the commitment of the associated mRNA substrates to NMD (see Introduction), we sought to elucidate these issues by combining yeast genetics with selective ribosome profiling and co-sedimentation analyses of wild-type and mutant Upfs. The results of these experiments strongly support the notion that all three Upfs manifest NMD functions while associated with ribosomes. In support of this conclusion we have shown that: 1) some *upf1* mutations that inhibit NMD also alter the extent to which Upf1 co-sediments with large polyribosomes (Figures 1 and 2), an observation noted previously with different *upf1* constructs^9^; 2) deletion of the *UPF2* gene markedly increases the extent to which Upf1 associates with polyribosomes, as well as the size of the polysomes with which Upf1 co-sediments (Figure 2a); 3) deletion of *UPF1* leads to co-sedimentation of Upf2 with polysomes that are lighter than those with which it co-sediments in WT cells (Figure 2a); 4) the extent of Dcp2 co-sedimentation with large polysomes is influenced by its interaction with Upf1 (Figure 2b); 5) immunopurification of ribosomes respectively bound by Upf1, Upf2, or Upf3 led to stochiometric recovery of each Upf factor with a large fraction of ribosomal proteins (Figure 3); and 6) Upf1 association with ribosomes in the presence of CHX leads to formation of an atypical footprint derived from the combined RNase1 protection of the mRNA by the ribosome and bound Upf1 (Figures 5 and 6). Finally, we note that ribosomes purified by virtue of their association with any of the three Upf factors protect mRNA segments enriched for NMD substrates (Figure 4a), i.e., these samples include ribosomes that are engaged in NMD or otherwise committed to the pathway. We conclude that, at least in yeast, Upf1, Upf2, and Upf3 are not displaced by elongating ribosomes, but are carried by them as they monitor translation termination and target aberrantly terminating mRNAs for NMD.

### Association of Upf1 with Ribosomes in CHX-treated Cells Promotes the Formation of a Ribosome:Upf1 Complex That Comprises an Early Step in NMD

In CHX-treated cells, the footprints recovered from immunopurified Upf1-associated ribosomes include the expected^37^ M fragments of ∼27-32 nt as well as novel 37-43 nt ribosome-protected fragments that we designate as L footprints (Figure 5a). The latter footprints may be formed by a CHX-induced ribosome collision and subsequent endonucleolytic cleavage^39^ or by the combined mRNA protection effects of Upf1 plus a ribosome to which it is bound (Figure 6). The latter additive effect would be consistent with the size of ribosome footprints and the observation that human Upf1 protects 8-11 nt of RNA in RNA binding and unwinding assays^40^. L footprints accumulate early during translation of mRNA ORFs, well before distal regions of mRNA are recovered in M or S footprints from Upf1-, Upf2-, or Upf3-associated ribosomes (Figure 7). L footprints are not recovered from Upf1-associated ribosomes purified from cells without CHX treatment (Figure 5a), nor are they derived from Upf2- or Upf3-associated ribosomes purified from cells with or without CHX treatment (Figure 5a), and their formation does not depend on specific placement of the FLAG epitope tag to one end of Upf1 (Figure 5). Formation of L footprints does depend on Upf1 function because they are absent when Upf1 harbors DE572AA substitutions, but they are not dependent on a functional NMD pathway or Upf2 because they are still recovered from Upf1-associated ribosomes in CHX-treated *upf2Δ* cells (Figure 5b and Extended Data Figure 6b). This combination of properties suggests that CHX treatment has trapped ribosomes and Upf1 in an otherwise transient state preceding a commitment to NMD, i.e., L footprints are Upf1- specific, not NMD-specific. The existence of such a transient state implies that Upf1 can interact with ribosomes in modes that either do or do not lead to NMD. The early appearance of L footprints (i.e., their recovery from 5’ regions of mRNA ORFs), and their persistence throughout most of the coding region (Figure 7, Extended Data Figure 6), implies that this potential surveillance for functional targets (e.g., a ribosome undergoing premature termination) is likely to be active during most translation elongation events. In addition, the uniformly delayed recovery of S and M footprints from Upf1-, Upf2, or Upf3-associated ribosomes (Figure 7, Extended Data Figure 6) indicates that: 1) on a transcriptome-wide basis most endogenous NMD substrates do not undergo premature or otherwise inefficient termination until at least 10-15% of the mRNA coding region has been translated and 2) once Upf1 commits to a substrate Upf2 and Upf3 exert their influence shortly afterwards.

### Preferential Association of NMD Substrates with Ribosomes Harboring Bound Upf Factors: Implications for the NMD Pathway

Our model for yeast NMD posits that all three Upfs stably associate with the ribosome during a termination event that is premature or otherwise inefficient and they should thus manifest a preference for association with ribosomes translating NMD substrates^1, 41^. Here, using selective ribosome profiling, we have demonstrated that Upfs are indeed preferentially bound to ribosomes engaged in translating NMD substrates and reduced in their association with non-NMD substrates (Figure 4a). These results support our model and confirm our earlier comparable result for Upf1 that was obtained using a completely different approach^35^. The general recovery of a large fraction of non-NMD substrates in our immunopurification procedure implies that all three Upf factors most likely interact stochastically with ribosomes, a step consistent with an mRNA surveillance function, or that the recovery of non-NMD substrates is a consequence of overexpression of the respective Upf factors. In either case, this result indicates that ribosome association is not the triggering event in NMD. The relative content of NMD substrates and co-purified proteins in different IP samples provides several insights into the sequence of events in the NMD pathway. The co-purification of Upf1 in Upf2 IP samples but not vice versa (Figure 3) implies, not surprisingly, that Upf1:Upf2 complex formation on the ribosome occurs subsequent to Upf1 association with the ribosome and represents a step toward NMD commitment. The latter conclusion is reinforced by the relative loss of NMD substrate association with Upf1 IP samples in *upf2Δ* cells (Figure 4b and Extended Data Figure 3b).

We reported recently that NMD substrates were poorly translated mRNAs, i.e., they had significantly lower ribosomal density throughout their open reading frames than non-NMD substrates, and that this diminished translational efficiency was not attributable to previously postulated translational repression effects of Upf1^36^. Here, we have confirmed these results, and have done so with a much larger sampling of strains and growth conditions than considered previously. Combining our previous analyses (based on published datasets) with the 13 datasets presented here (Figure 4c) strongly supports the notion that NMD substrates are translated inefficiently, raising the question of the basis of this property. We noted previously that codon optimality was lower for NMD substrates than for non-substrates^36^, but another feature of NMD substrates that may affect their translation is the possible preponderance of upstream open reading frames (uORFs) in these mRNAs. The uORFs in question would include those that are bona fide regulators of specific gene expression, as well as those arising as a consequence of indiscriminate transcription^42^. The latter may also account for the larger isoforms of NMD substrates we had identified previously^35^.

Finally, there is the question of how polyribosomes with associated Upf proteins become targets for accelerated mRNA decay. NMD in yeast initiates with mRNA decapping^43^ while mRNAs are polysome-associated^44^ and this process is dependent on Dcp2, the catalytic component of the decapping enzyme. We have shown recently that Dcp2 contains two nearly identical binding sites for Upf1^27, 28^ and that Upf1:Dcp2 association controls the selective targeting of the decapping enzyme to NMD substrates.^28^ The ability of Upf1 to dimerize^24^, and the presence of two Upf1 binding sites on Dcp2, suggests that Dcp2 may be recruited to polyribosomes by dimerization of ribosome-associated Upf1 with Dcp2-associated Upf1.

## METHODS

### Yeast Strains

Yeast strains used in this study are listed in Supplementary Table 3. Strains containing complete gene deletions of *UPF1*, *UPF2*, and *UPF3* (HFY871, HFY115, HFY861, HFY467) were described previously^25, 45^. HFY2996 and HFY3039 were described previously^28^. YRG5205 was generated from HFY3039 with a complete gene deletion of *UPF1* as in HFY871.

### Oligonucleotides

Oligonucleotides used in this study were obtained from Eurofins Operon or Integrated DNA Technologies (IDT) and are listed in Supplementary Table 4. Gene fragments were synthesized by Quintara Biosciences and are described below in Plasmids.

### Plasmids

Plasmids used in this study are listed in Supplementary Table 5. The following plasmids were published previously: YEplac112^46^; pG1-*FLAG-UPF1*^47^; YEplac112-*FLAG-UPF2* (also known as YEplac112-F2-*NMD2*)^25^; pRS425 and pRS316^48^; pRS315-*NMD2*-C-Sal^49^; YEplac195-*TPI*-*FLAG-UPF3*^50^.

YEplac112-*6XHis-UPF1*: pRS314-*UPF1*^25^ was digested with BamHI, SalI, and XbaI to release a 4.2kb BamHI-SalI fragment of *UPF1*. This fragment was ligated into pGEM-3Zf (+) that had been digested with BamHI and SalI and dephosphorylated with calf intestinal alkaline phosphatase (NEB). 6XHis tag was added to the N-terminus of Upf1 in pGEM-3Zf (+) using QuickChange Lightning Site-Directed Mutagenesis Kit (Agilent) with oligonucleotides 5-N-His-UPF1 and 5-N-HIS-UPF1-r to yield pGEM3Zf(+)-*6XHis*-*UPF1*. A ∼4.2kb SacI/PstI fragment was isolated from pGEM3Zf(+)-*6XHis*-*UPF1*, ligated into YEplac112 which had been digested with SacI and PstI and dephosphorylated with calf intestinal alkaline phosphatase to yield YEplac112-*6XHis-UPF1*.

pRS425-*UPF1-FLAG*: gene fragments were synthesized by Quintarabio with flanking restriction sites 1) UPF1-Xho-Sph, a 1593 bp fragment containing the *TDH3* (glyceraldehyde-3-phosphate dehydrogenase, GAPDH) promoter inserted into a BamHI site plus ggc (ggatccggc) immediately upstream of the start codon of the *UPF1* coding region, then from the start codon to the SphI site at position 914 of the *UPF1* coding sequence ; 2) UPF1-Sph-Nco, a 1102 bp fragment from the SphI site in the *UPF1* coding sequence to the NcoI site in the *UPF1* coding sequence; 3) UPF1-Nco-Sac, a 1167 bp fragment from the NcoI site in the *UPF1* coding sequence to a SacI site in the *UPF1* 3’UTR followed by a spacer with a SacI restriction site (caccgcggtggagctc), and FLAG epitope sequence (gattacaaggatgacgacgataag) inserted immediately upstream of the stop codon. These three gene fragments were ligated into pRS425 at the XhoI-SacI sites of the MCS.

Mutations of *UPF1* were placed into *UPF1-FLAG* as follows:

*<u>C62Y</u>*: The UPF1-Sph-Nco and UPF1-Nco-Sac gene fragments described above were cloned in to the SphI-SacI sites in the pGEMT-EZ (Promega) MCS to generate pGEMT-EZ-Sph-*UPF1-FLAG*-Sac.The UPF1-Xho-Sph gene fragment in plasmid vector pQ (Quintarabio) was mutagenized with oligonucleotides 5Upf1C62Ymut and 3Upf1C62Ymut using the QuikChange II XL site directed mutagenesis kit (Agilent). The UPF1XhoI-C62Y-SphI fragment was isolated and ligated with the 2.28 kb Sph-*UPF1-FLAG*-Sac fragment from pGEMT-EZ-Sph-*UPF1-FLAG*-Sac into the XhoI-SacI site of the MCS of pRS425.

*<u>K436E</u>*: pGEM3Z-6XHis-*UPF1* was mutagenized with oligonucleotides K436F and K436R using the QuikChange II XL site directed mutagenesis kit, the 1116 bp SphI-NcoI fragment containing the *K436E* mutation was isolated and substituted for the UPF1-Sph-Nco fragment as described above for the construction of pRS425-*UPF1-FLAG*.

*<u>AKS484HPA</u>*<u>, *RR793AA* and *R779C*</u>: pGEMT-EZ-Sph-*UPF1-FLAG*-Sac was mutagenized with oligonucleotides Upf1AKS-HPAmut and UPF1AKS-HPAmutrev, 5Upf1RRAAmut and 3Upf1RRAAmut, or Upf1R779C and Upf1R779Crev, respectively, using the QuikChange II XL site directed mutagenesis kit. The 2.28 kb SphI-SacI fragments were isolated and were ligated with the UPF1-Xho-Sph gene fragment into the XhoI-SacI site of the pRS425 MCS.

*<u>DE572AA</u>*: the 1116bp SphI-NcoI fragment from pACTII-*UPF1*-TH4-3-2-DE572AA ^24^ was isolated and substituted for the UPF1-Sph-Nco fragment described above.

*UPF1-FLAG* and mutants were placed under the *UPF1* promoter into pRS316 as follows: a 340 bp fragment containing the *UPF1* promoter sequence flanked by XhoI and BamHI restriction sites was generated by PCR from pRS313-*UPF1* (E-B) with oligonucleotides 5Upf1XhoProm and 3Upf1BamProm and digested with XhoI and BamHI. A 3.195 kb BamHI-SacI fragment from pRS425-*UPF1-FLAG* or mutants was isolated and ligated with the digested PCR product above into the XhoI-SacI site of the pRS316 MCS.

pRS425-*UPF2-FLAG*: gene fragments were synthesized by Quintarabio with flanking restriction sites 1) UPF2-Xho-Bam, from an XhoI site inserted 547nt upstream of the start codon, including the intron, to the BamHI site at position 537 of the coding sequence; 2) UPF2-Bam-RI from the BamHI site to the EcoRI site at position 2314 of the coding sequence; 3) UPF2 RI-Sac, from the EcoRI site to a SacI site inserted 500nt downstream of the stop codon, and the FLAG epitope sequence inserted immediately upstream of the stop codon. These three gene fragments were ligated in to pRS425 at the XhoI-SacI sites of the MCS.

pRS425-*UPF2*: a 4946 bp fragment generated by PCR with primers Upf2-425-fwd and Upf2-425-rev containing a the entire UPF2 gene was inserted in to pRS425 cut with NotI and SalI using the NEBuilder High-Fidelity DNA Assembly Cloning Kit (New England Biolabs).

YEplac195-*TPI-UPF3*: A 1164 bp SalI-XbaI fragment was generated by PCR from genomic DNA with primers 5’Sal-UPF3 and 3’Xba-UPF3 and restriction digested with SalI and XbaI. YEplac195-*TPI-FLAG-UPF3* was digested with SalI and XbaI and the vector backbone was ligated to the digested PCR product to generate YEplac195-*TPI-UPF3*.

pRS425-*FLAG-UPF1*: a 667 bp XhoI-BamHI fragment from the UPF1-Xho-Sph gene fragment, a 926 bp BamHI-SphI fragment from pG1-*FLAG-UPF1*, and the UPF1-Sph-Nco and UPF1-Nco-Sac gene fragments were ligated in to pRS425 at the XhoI-Sac1 sites of the MCS.

### Polyribosome analysis, protein detection, and quantitation

For polyribosome analysis, cells were grown in selective media and harvested as described previously ^10^. Lysates were loaded onto 7-47% sucrose gradients, subjected to ultracentrifugation, fractions collected, and individual fractions were precipitated with trichloroacetic acid (TCA). Samples from total or immunopurified ribosomes or aliquots from TCA precipitated sucrose gradient fractions were run in 1X Laemmli buffer on 4-20% Mini-PROTEAN TGX Precast Protein Gels (BioRad) in 1X Tris/Glycine/SDS buffer (BioRad), subjected to western blotting onto Immobilon™-P PVDF Transfer Membranes (EMD Millipore) by electrotransfer using a Trans-Blot® SD Semi-Dry Transfer Cell (Biorad), and protein was visualized with rabbit α-FLAG polyclonal, mouse α-HA monoclonal antibody (Sigma) or rabbit monoclonal α-L3 (Tcm1) (gift from Dr. Jon Warner) followed by HRP-conjugated goat α-mouse (Life technologies) or donkey α-rabbit (GE) antibody. Polysomal distribution of tagged WT and mutant Upf1, Upf2 or Dcp2 protein and tagged protein signal from total or immunopurified ribosomes was visualized by western blotting on Amersham Hyperfilm ECL film (GE). Band quantitation from scanned images was performed using Multigauge V3.0 (Fuji). For analysis of polysome distribution of tagged proteins, the protein signal for a given fraction was calculated as the percent of its total signal across fractions 1-10 of an individual sucrose gradient western blot. Values from western blots of replicate gradient fractions were averaged and heatmaps generated to display the average percent signal across sucrose gradient fractions 1-10 from all replicates of that sample.

### Ribosome purification

Yeast cultures were grown, whole cell yeast lysates were prepared and digested with RNaseI, and ribosomes were recovered as described previously^30^.

### Immunopurification

FLAG-tagged protein-associated ribosomes were isolated by immunopurification using anti-FLAG M2 Affinity gel as described previously^30^. Eluate was collected by centrifugation and spun through Amicon Ultra 2ml 100K centrifugal filter units (Millipore) until the volume was <200μl. The concentrate from one reaction set was pooled into one Amicon Ultra 2ml 100K centrifugal filter unit and spun until the total volume was approximately 160μl. For the *UPF1-FLAG*, *DE572AA-FLAG* and *UPF1-FLAG/upf2Δ* ribosome profiling libraries, immunopurified ribosomes were flash frozen and stored at -80°C until RNA purification. For all other ribosome profiling libraries and for all mass spectrometry analyses, the concentrate was layered on top of a 1M sucrose cushion in IP buffer plus 1X protease inhibitors, 10U/ml Superase-In, and spun at 166,180g for 100 minutes at 4°C in a TLA100 rotor. Supernatant was removed and the pellet was resuspended in 60μl IP buffer plus 0.5mM DTT, 1X protease inhibitors, 20U/ml Superase-In; and A_260_ was determined on a spectrophotometer.

### Sample Preparation for Mass Spectrometry

Mass spectrometry analyses were performed by the Mass Spectrometry Facility of the UMass Chan Medical School. Samples were prepared for mass spectrometry by disrupting total or immunopurified ribosomes in 1X Laemmli buffer and running on a 4-20% Mini-PROTEAN TGX Precast Protein Gel in 1X Tris/Glycine/SDS buffer (BioRad) for 5 minutes. Gels were stained using the Colloidal Blue Staining Kit (BioRad) and destained in water. Fragments were excised and cut into 1x1 mm pieces, transferred to microfuge tubes, and each added 1 mL of water, followed by a solution of 20 µL of 45 mM of 1,4 dithiothreitol (DTT) in 200 µL of 250 mM ammonium bicarbonate. Samples were incubated at 50°C for 30 minutes, cooled to room temperature, added 20 µL of 100 mM iodoacetamide (IAA) and incubated for 30 min. Excessive DTT and IAA was removed, and the gel pieces washed with water (3x, 1 mL each), followed by 1mL of a 1:1 solution of 50 mM ammonium bicarbonate:acetonitrile, quenched with 200 µL of acetonitrile and dried in a Speed Vac. Gel pieces were rehydrated in a mixture of 4 ng/µL trypsin (Promega, Madison, WI) and 0.01% ProteaseMAX (Promega) in 50 µL of 50 mM ammonium bicarbonate and incubated for 18 hours at 37°C. Supernatants were collected and further extraction was performed by adding 200 µL of 80:20 solution of acetonitrile:1% (v/v) formic acid in water. Supernatants were combined, peptides were lyophilized in a Speed Vac and resuspended in 25 µL of 5% acetonitrile with 0.1% (v/v) formic acid for mass spectrometry analysis.

### Mass Spectrometry

Data was acquired using a NanoAcquity UPLC (Waters Corporation, Milford, MA) coupled to a Q Exactive hybrid mass spectrometer (Thermo) (N-terminal FLAG-tagged datasets) or an Orbitrap Fusion Lumos Tribrid (Thermo Fisher Scientific, Waltham, MA) mass spectrometer (C-terminal FLAG-tagged datasets). Peptides were trapped and separated using an in-house 100 µm I.D. fused-silica precolumn (Kasil frit) packed with 2 cm ProntoSil (Bischoff Chromatography, DE) C18AQ (200Å, 5µm) media and configured to an in-house packed 75 µm I.D. fused-silica analytical column (gravity-pulled tip) packed with 25 cm Magic (Bruker, Billerica, MA) C18AQ (100Å, 3µm) media, respectively. Mobile phase A was 0.1 % (v/v) formic acid in water and mobile phase B was 0.1 % (v/v) formic acid in acetonitrile. Following a 3.8 µL sample injection, peptides were trapped at flow rate of 4 µL/min with 5% B for 4 min, followed by gradient elution at a flow rate of 300 nL/min from 5-35% B in 60 min, wash and reconditioning (total run time ∼90 min). Electrospray voltage was delivered by liquid junction electrode (1.4 kV) located between the columns and the transfer capillary to the mass spectrometer was maintained at 275°C. Mass spectra were acquired over *m/z* 300-1750 Da with a resolution of 60,000 (*m/z* 200), maximum injection time of 30 ms, and an AGC target of 700,000. Tandem mass spectra were acquired using data-dependent acquisition (2 sec cycle) with an isolation width of 1.2 Da, HCD collision energy of 30%, resolution of 15,000, maximum injection time of 100 ms, and an AGC target of 10,000. Biological triplicates of each strain (input and IP) were analyzed in technical triplicate.

### Database Searches

Raw data were processed using Proteome Discoverer (Thermo, version 2.1.1.21) and searched against Uniprot Yeast (downloaded 07/2021) database using Mascot (Matrix Science, version 2.6.2). Search parameters were as follows: full tryptic specificity with up to 2 missed cleavages; precursor mass tolerance 10 ppm; fragment mass tolerance 0.05 Da; cysteine carbamidomethylation considered as a fixed modification, while protein N-terminal acetylation, methionine oxidation, peptide N-terminal (E,Q) pyroglutamate conversion, serine/threonine phosphorylation and lysine ubiquitination (GG) were specified as variable modifications. Peptide and protein validation and annotation was done in Scaffold 4.8.9 (Proteome Software, Portland, OR) employing Peptide Prophet ^51^ and Protein Prophet ^52^ algorithms. Peptides were filtered at a 1% FDR, while protein identification threshold was set to greater than 99% probability and with a minimum of 2 identified peptides per protein. Intensity-based absolute quantification (iBAQ) ^31, 32^, calculated and normalized across biosamples in Scaffold, is used as a measure of protein abundance. For each protein in a sample, at least two replicates were required to have positive iBAQ values for averaging and plotting; otherwise, the identification of the protein was deemed inconsistent and average iBAQ was set to 0.

### mRNA Decay Analysis

mRNA decay phenotypes were assessed by northern blotting of total RNA prepared from cell pellets by the hot phenol method^53^, probing for *CYH2* mRNA and pre-mRNA as described previously^8, 25, 53^. Blots were visualized on a Fuji phosphorimager and quantitated with Multigauge V3.0 (Fuji). Northern blot images in Figure 1a were subjected to a Multigauge noise reduction filter, kernel size 3X3, applied equally across the entire image for display purposes only.

### RNAseq and Ribosome Profiling Library Preparation from Cells Expressing N-terminal FLAG-tagged Upf Proteins

RNA was isolated from clarified lysates (60 μl), and total (15 μl) or immunopurified ribosomes (entire recovered volume) using the miRNeasy mini kit (Qiagen) according to manufacturer’s standard protocol. RNA was eluted in 30 μl (lysate) or 14 μl (ribosomes), then treated with RiboZero Magnetic Gold (Yeast) Kit (Illumina) as described previously^30^. Ribosome protected RNA fragments were 3’ dephosphorylated with T4 polynucleotide kinase (NEB) in 150 mM MES-NaOH pH 5.5, 450 mM NaCl, 15 mM MgCl2, 15 mM β-mercaptoethanol, 0.3U/μl Superase-In at 37°C, 2 hours; 65°C, 20min; and RNA was selectively recovered with RNA clean and concentrator-5 (Zymo Research). RNA fragments were then 5’ phosphorylated with T4 polynucleotide kinase in T4 PNK buffer, 1mM ATP, 1U/μl Superase-In at 37°C, 1 hour; 60°C, 10 min; and RNA was selectively recovered with RNA Clean and Concentrator-5 (Zymo) according to the manufacturer’s protocol. Ribosome profiling libraries were prepared using the NEXTflex Small RNA-Seq Kit v3 (Perkin Elmer).

Total mRNA libraries were prepared using the TruSeq Stranded mRNA LT Sample Prep Kit (Illumina) according to manufacturer’s protocol.

Over the course of library preparation, the amounts of RNA and final libraries were quantified by Qubit 3.0 Fluorometer with the Qubit RNA HS Assay Kit (Invitrogen) and the Qubit dsDNA HS Assay Kit (Invitrogen), respectively. Assessments of rRNA depletion, RNA quality, and final libraries were done by Fragment Analyzer capillary electrophoresis system (Advanced Analytical) at the UMass Chan Medical School Molecular Biology Core Labs (RRID: SCR_018263).

### RNAseq and Ribosome Profiling Library Preparation from Cells Expressing C-terminal FLAG-tagged Upf Proteins

Because the RiboZero Magnetic Gold (Yeast) Kit (Illumina) was discontinued midway through this study, we developed an alternative rRNA depletion method based on oligonucleotide blocking of adapter ligation and cDNA synthesis reactions. To maximize the amount of ribosome protected mRNA fragments input into the libraries, we altered immunoprecipitation and RNA isolation protocols for the C-terminally FLAG tagged ribosome profiling libraries as follows by: 1) eliminating the final pelleting through a sucrose cushion following immunopurification, to avoid any loss of immunopurified ribosomes, and 2) using the small (<200nt) RNA preparation workflow in the miRNeasy kit according to manufacturer’s instructions, to remove as much extraneous RNA (either rRNA or large mRNA fragments) as possible prior to ribosome profiling library preparation.

Following RNA isolation, 10 μl RNA from total or immunopurified ribosomes were 3’ dephosphorylated and 5’ phosphorylated as described above ^30^ and was selectively recovered with RNA Clean and Concentrator-5 (Zymo) according to the manufacturer’s instructions, using adjusted RNA Binding Buffer diluted 1:2 with ethanol, into 13 μl water. 8.5 μl RNA was incubated in a thermocycler with a preheated lid with 2 μl oligonucleotides (for RNA from total ribosomes) or 1 μl 1:10 diluted oligonucleotides (for RNA from immunopurified ribosomes) from the QIAseq FastSelect –rRNA Yeast Kit (Qiagen) in 10.5 μl total at 75°C, 2 min; 70°C, 2 min; 65°C, 2 min; 60°C, 2 min; 55°C, 2 min; 37°C, 2 min; 25°C, 2 min; 4°C, hold. Ribosome profiling libraries were prepared using the NEXTflex Small RNA-Seq Kit v3 (Perkin Elmer) but eliminating the 70 °C denaturation step prior to 3’ 4N Adenylated Adapter ligation. Adapters were undiluted for RNA from total ribosomes and 1:4 diluted for RNA from immunopurified ribosomes. PCR cycles were performed until a library appeared by Fragment Analyzer analysis, 15 cycles for libraries from total ribosomes or 15-18 cycles for libraries from immunopurified ribosomes, and libraries were selectively recovered using the gel-free size selection and cleanup protocol according to manufacturer’s protocol.

For RNAseq library preparation, RNAs from clarified lysates (60 μl) were isolated using the miRNeasy mini kit (Qiagen) according to manufacturer’s standard protocol. The use of TruSeq Stranded mRNA LT Sample Prep Kit (Illumina) was altered as follows: 1μg of RNA, 14.5 μl of Fragment, Prime, Finish mix (FPF), 1μl of oligonucleotides from the QIAseq FastSelect –rRNA Yeast Kit (Qiagen) in a total of 20.5 μl were incubated in a thermocycler with a preheated lid at 94°C, 8 min; 75°C, 2 min; 70°C, 2 min; 65°C, 2 min; 60°C, 2 min; 55°C, 2 min; 37°C, 2 min; 25°C, 2 min; 4°C, hold; followed by first strand cDNA synthesis and the remainder of the protocol according to manufacturer’s instructions, except RNA Adapters were diluted 1:2 at the ligation step. If excess adapter dimers were present, it was necessary to reconstruct the library and dilute the RNA Adapter 1:4 or 1:8 and increase the PCR cycle number to 17, as was done for libraries T5019, T5033, T5035 and T5338.

### High-throughput Sequencing of RNAseq and Ribosome Profiling Libraries

RNAseq libraries were sequenced on either a HiSeq4000 (Illumina) at Beijing Genomics Institute (BGI) or a NextSeq500 (Illumina) in-house. Ribosome profiling libraries were sequenced on a NextSeq500 in-house.

### Sequence Alignment, Transcript Quantification, and Data Analyses

RNA-seq and ribosome profiling reads were aligned to a yeast transcriptome (available at https://github.com/Jacobson-Lab/yeast_transcriptome_v5) using bowtie^54^ and transcript abundance were determined using RSEM^55^, as described previously^56^. For intron-containing genes, spliced (“mRNA”) and unspliced (“pre-mRNA”) isoforms were indexed as separate entries.

Ribosome profiling libraries were pre-processed with adapter trimming and removal of reads aligned to non-protein coding RNAs. Because some of the constructs were tagged at their N-termini, a fraction of reads recovered were from ribosomes associated with the Upf factor nascent peptide (Supplementary Table 6). Therefore, reads arising from *UPF* factor transcripts were discarded from these and all subsequent ribosome profiling libraries analyzed. PCR duplicates identified via the 4N barcodes on either side of the read were removed. Read processing statistics for ribosome profiling libraries are provided in Supplementary Table 6.

The remaining unique reads were further processed in two ways:

1. Transcript abundance was determined by RSEM either with all reads or a subset of reads of desired read lengths using a transcriptome containing pre-mRNA entries (rsem-calculate-expression --strandedness forward --fragment-length-mean [vary] -- fragment-length-sd [vary] --seed-length 15 --bowtie-m 10). Reproducibility of biological replicate libraries is shown as correlation matrix (Extended Data Figure 7, top) and PCA plots (Extended Data Figure 7, bottom).
2. Reads were aligned to a separate transcriptome where only spliced mRNAs were considered (bowtie -m 10 -n 2 -l 15). The resulting aligned reads were processed by R package riboWaltz^57^ for initial visual inspection and calculation of P-site offsets, which were manually checked and modified for accuracy (Supplementary Table 7) and used as the basis for subsequent reading frame calculations, periodicity, and other metagene plots. Analysis of ribosome footprints from both CHX treated and untreated cells and from total and Upf factor associated ribosomes showed the footprints mapped primarily to coding regions, most P-sites mapped to the “0” reading frame in coding regions, and displayed 3-nucleotide periodicity, indicative of translating ribosomes (Extended Data Figures 5, 6, 8 and 9). As expected, CHX addition resulted in accumulation of ∼27-32 nt footprints over the start codon, as described previously^38, 58, 59^. Replicates were averaged unless otherwise specified.

RNA-seq libraries were aligned to the transcriptome containing pre-mRNA entries and transcript abundance determined by RSEM (rsem-calculate-expression --strandedness reverse --fragment-length-mean 200 --fragment-length-sd 50 --bowtie-m 10) without any pre-processing steps. Reproducibility of biological replicate RNAseq libraries is shown as a correlation matrix (Extended Data Figure 10a) and PCA plots (Extended Data Figure 10b).

### Analyses of Changes in Transcript Abundance

All analyses involving transcript abundance changes (translation efficiency: log_2_(Total ribo-seq / RNAseq); log_2_(IP / Total); log_2_(L / M); log_2_(M / S)) were performed in the R programming environment. The “expected_count” columns in the RSEM file output “isoforms.results” were used as input. An R package DESeq2 ^34^ was used to normalize read counts and calculate log_2_ fold changes in transcript abundance with the default low count and outlier filtering parameters. The median log_2_ fold changes belonging to NMD substrates was compared to non-NMD substrates using Wilcoxon’s rank sum test with Benjamini-Hochberg method for multiple testing correction. Adjusted p-values are reported. Standardized effect sizes were determined by wilcoxonRG() function from an R package rcompanion. Fisher’s exact test was performed by fisher.test() function and logistic regression by glm() function with “family” parameter = binomial(link = “logit”) in R stats package. R packages used for data preparation, statistical analyses, and plotting were DEseq2, riboWaltz, stats, rcompanion, ggplot2, ggpubr, ggrepel, ggh4x, data.table, dplyr, and reshape2.

### Data Availability

All custom scripts have been made available at https://github.com/Jacobson-Lab/Upf-polysomes. Sequencing data that support this study have been deposited in the National Center for Biotechnology Information Gene Expression Omnibus (GEO) with the accession code GSE186795 (https://www.ncbi.nlm.nih.gov/geo/query/acc.cgi?acc=GSE186795). The mass spectrometry proteomics data have been deposited to the ProteomeXchange Consortium via the PRIDE partner repository^60^ with the dataset identifier PXD029577 and 10.6019/PXD029577.

## ACKNOWLEDGMENTS

This work was supported by a grant to A.J. (1R35GM122468) from the U.S. National Institutes of Health. We thank Alper Celik for his generous gifts of yeast strains and plasmids; Chan Wu, Alper Celik, EiEi Min, Scott Shaffer, Andrei Korostelev, Zhiping Weng, Sean Ryder, John Leszyk, Michelle Dubuke, and Khaja Muneeruddin for helpful discussions; and the UMass Chan Medical School Mass Spectrometry Core Facility for extensive proteomics analyses. We also thank the late Dr. Jon Warner for his generous gift of α-L3 (Tcm1) antibody.

## AUTHOR CONTRIBUTIONS

R.G. and A.J. conceived and designed the experiments, R.G. and K.M. carried out the experiments, R.B. and K.M. wrote data processing scripts, K.M., R.B., R.G., F.H., and A.J. analyzed the data, R.G., K.M., and A.J. wrote the paper with input from all authors, and A.J. obtained funding for the study.

## ADDITIONAL INFORMATION

Supplementary Information is available for this paper.

Correspondence and requests for materials should be addressed to Allan Jacobson.

Reprints and permissions information are available at www.nature.com/reprints

## EXTENDED DATA

**Extended Data Figure 1.**
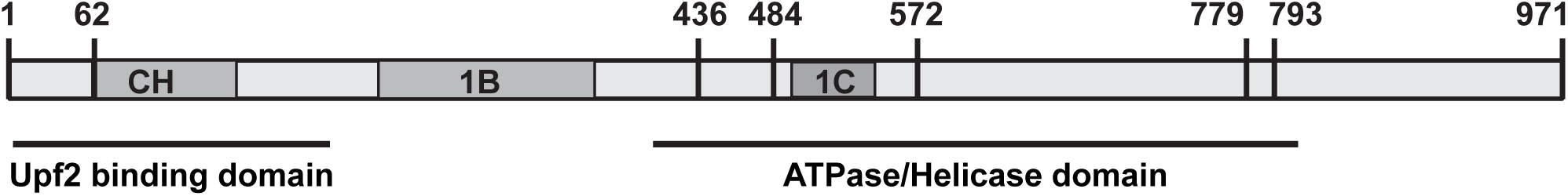
Schematic representation of Upf1 protein and domain structure. Numbers indicate amino acid location of alleles characterized previously ^9, 12, 22, 24, 61–63^. Upf2 binding, CH, 1B, 1C and ATPase/helicase domains^45, 61^ are indicated.

**Extended Data Figure 2.**
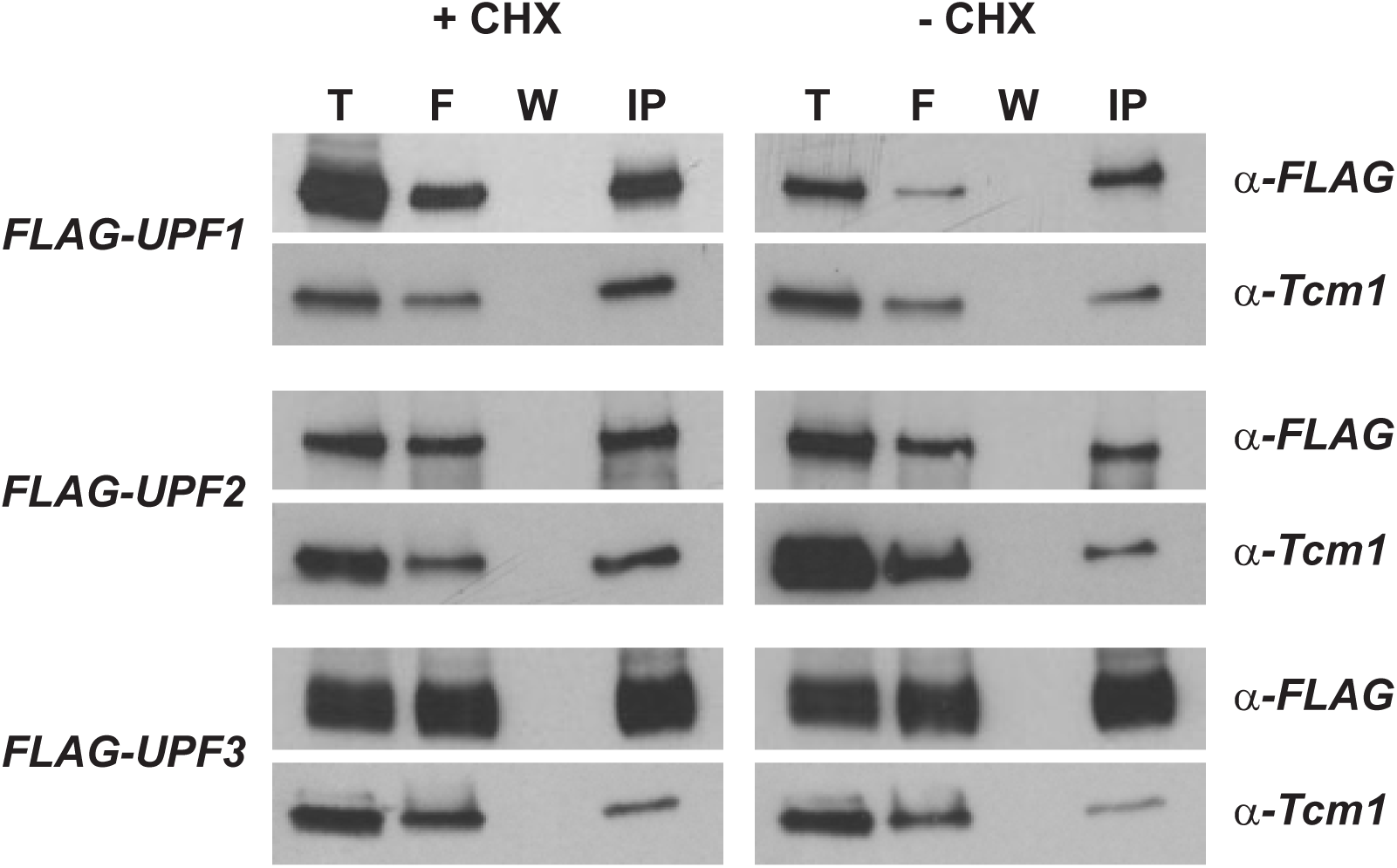
Immunopurification of ribosomes associated with FLAG-tagged Upf factors. Representative western blots of aliquots of total (T), flowthrough (F), final wash (W), and pelleted immunopurified ribosomes (IP) derived from strains expressing episomal *FLAG-UPF1*, *FLAG-UPF2,* or *FLAG-UPF3*. Lysates were prepared in the presence (+CHX) or absence of cycloheximide (-CHX) and ribosomes were immunopurified with α-FLAG resin. Blots were probed with α-FLAG antibody or α-Tcm1 antibody. These images were published previously^30^.

**Extended Data Figure 3.**
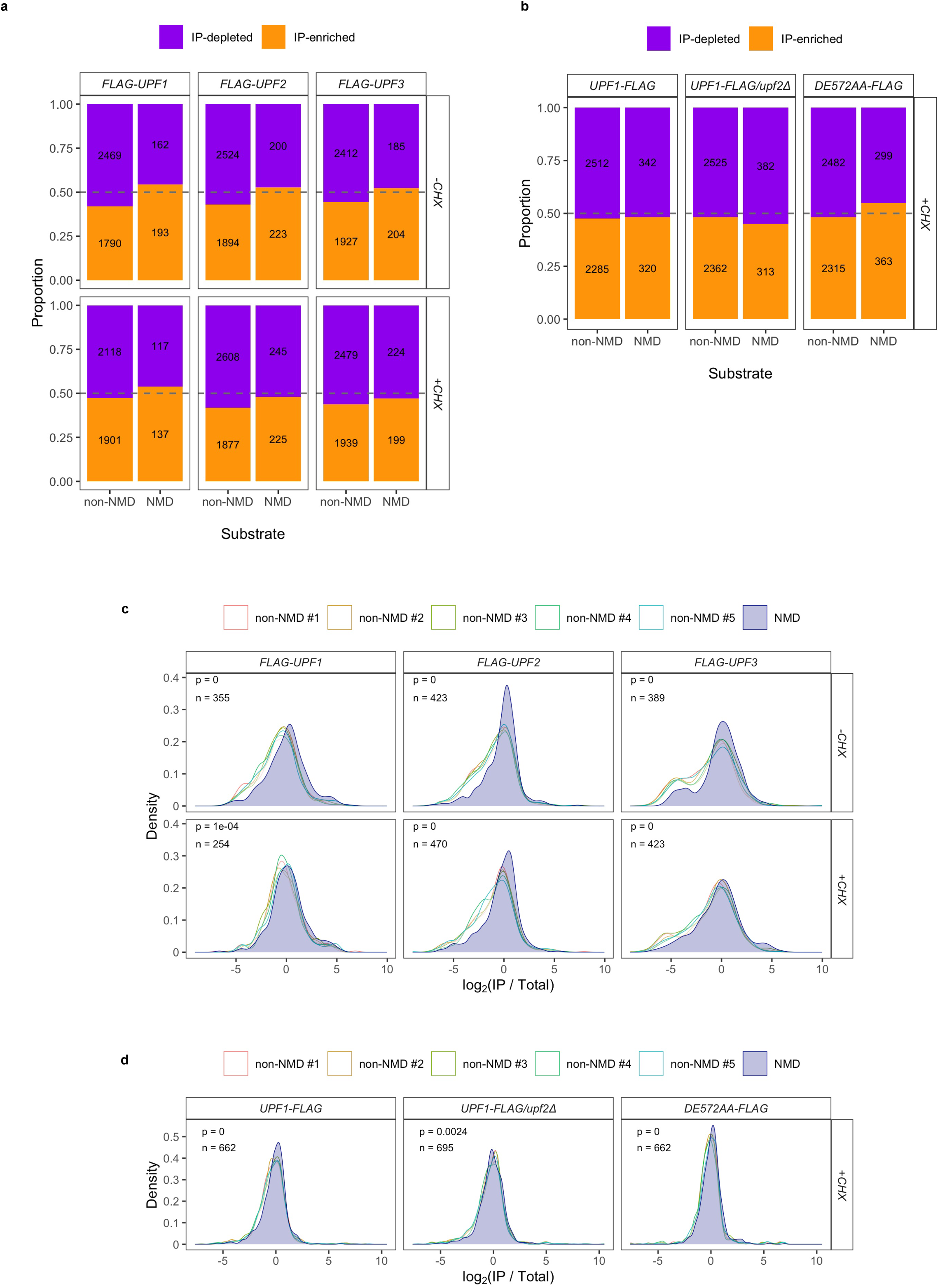
NMD substrates are enriched in Upf factor-associated ribosomes relative to total ribosomes. **a**-**b,** Proportions of NMD or non-NMD substrates in IP-enriched (log_2_(IP/Total) > 0, orange) and IP-depleted (log_2_(IP/Total) < 0, purple) were obtained and plotted. The number of transcripts representing each portion of the bar is reported. **c**-**d,** Analysis as in Figure 4 with additional constraints. Non-NMD substrates with average normalized read counts in IP and Total ribosomes matching those of NMD substrates were randomly subset 10,000 times, five of which were plotted. Empirical p-value was the frequency of when median log_2_(IP/Total) of non-substrates is as high as or higher than that of NMD substrates. **a** and **c,** Data from strains expressing *FLAG-UPF1*, *FLAG-UPF2,* or *FLAG-UPF3*. **b** and **d,** Data from strains expressing *UPF1-FLAG*, *UPF1-FLAG*/*upf2Δ*, or *upf1-DE572AA-FLAG*.

**Extended Data Figure 4.**
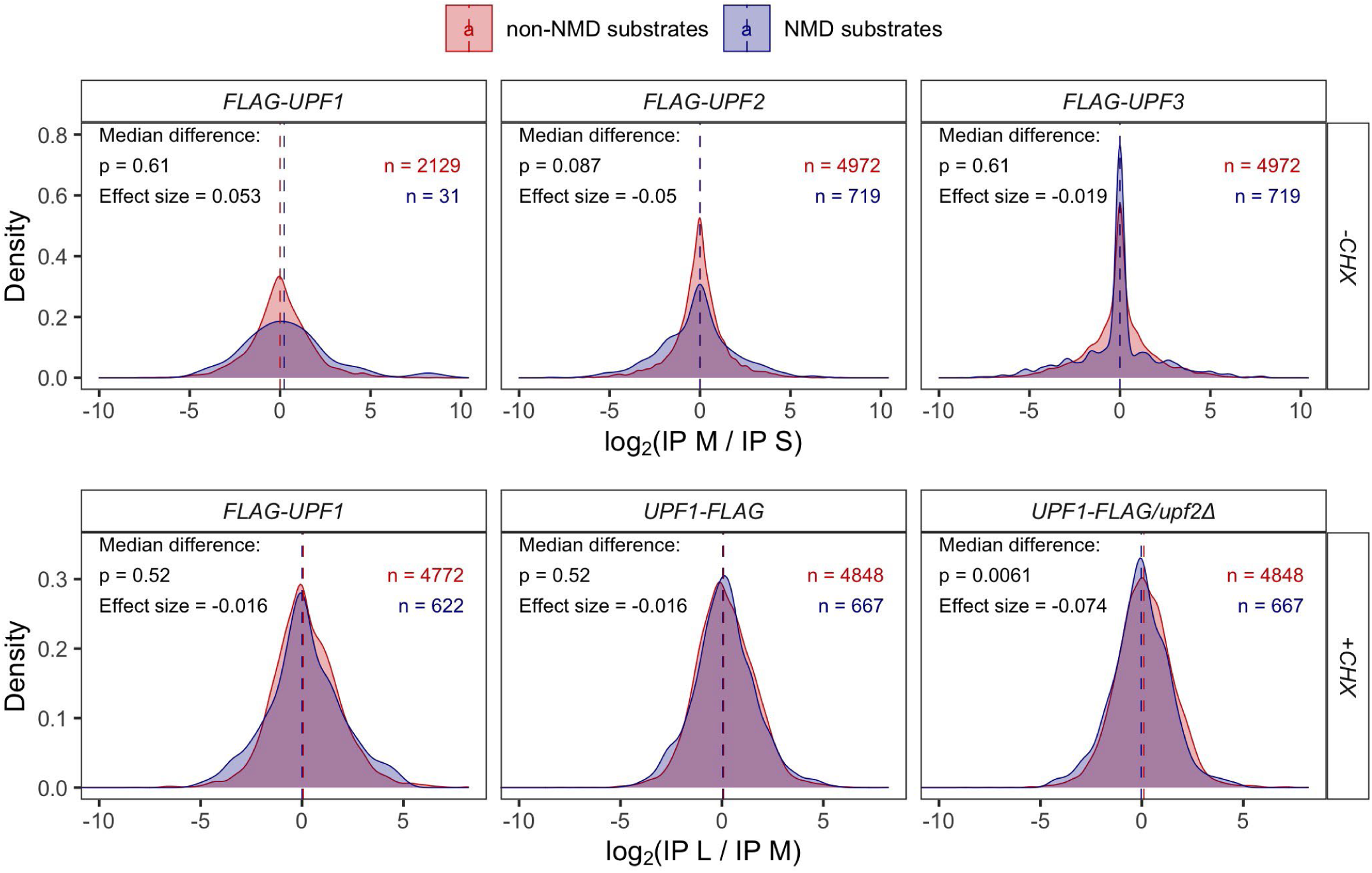
NMD substrate enrichment in Upf-associated ribosomes is not specific to particular footprint size class. For *FLAG-UPF1*, *FLAG-UPF2,* and *FLAG-UPF3* strains (-CHX), M and S footprints abundance from immunopurified (IP) ribosomes were normalized to the corresponding M and S footprint abundance from the input (Total) ribosomes. Thus, the relative change of M over S abundance was log_2_((IP M / Total M) / (IP S/ Total S)). For *FLAG-UPF1*, *UPF1-FLAG*, and *UPF1-FLAG/upf2Δ* strains (+CHX), L and M footprints abundance from IP were both normalized to the M footprint abundance from Total. The relative change of L over M was log_2_((IP L / Total M) / (IP M/ Total M)). Median log_2_ fold changes of minor and major footprint abundance of NMD substrates and those of non-substrates were compared using Wilcoxon’s rank sum test with Benjamini-Hochberg method for multiple testing correction. The effect size indicates the degree of difference between median log_2_ fold change of NMD substrates and that of non-substrates.

**Extended Data Figure 5.**
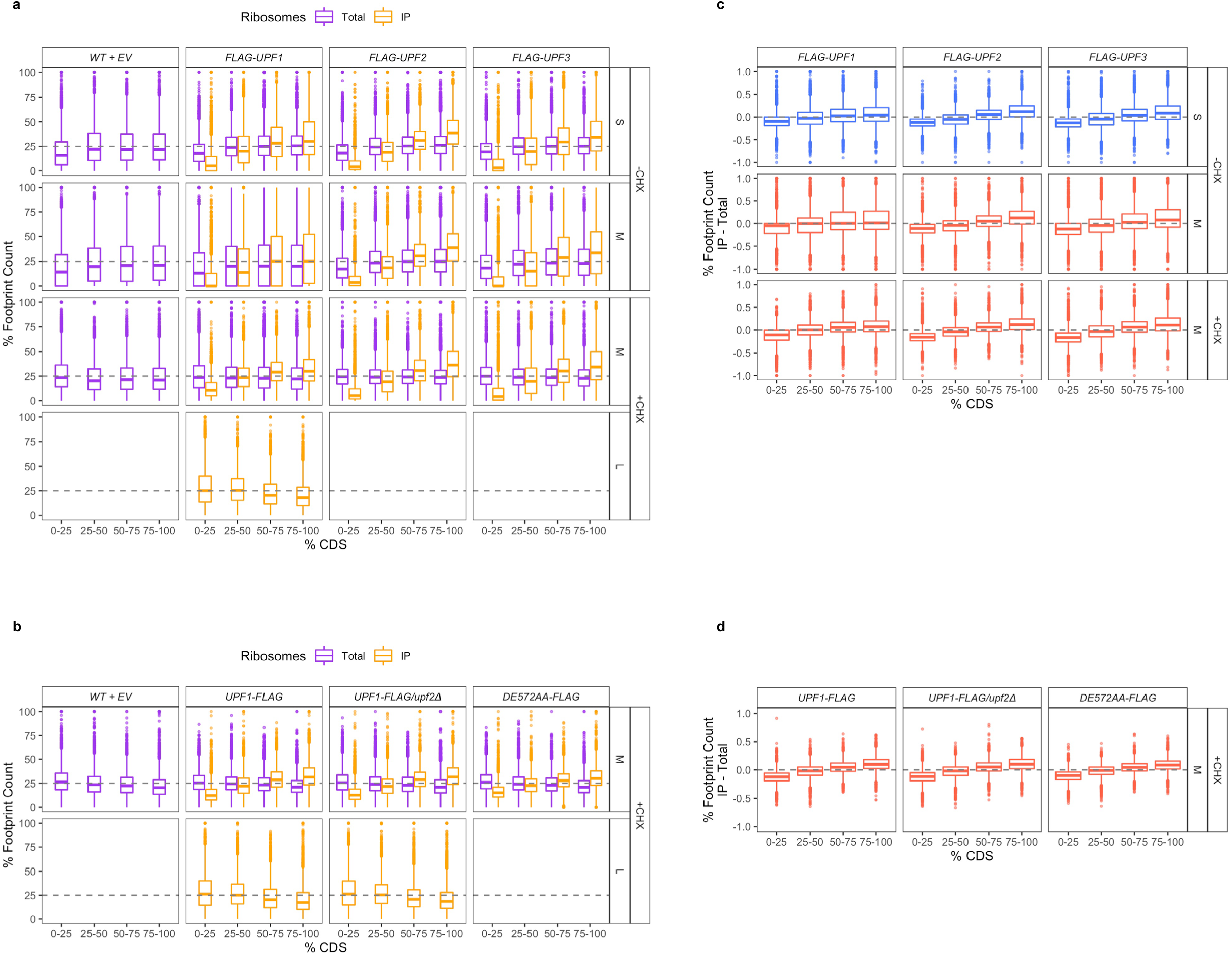
Upf factors progressively associate with ribosomes across mRNA coding regions except when Upf1 association forms L footprints. **a**-**b**, Distribution of footprint abundance across the coding region (CDS) for each footprint size class from total (purple) or IP (orange) ribosomes profiling libraries Each gene’s CDS was divided into 4 bins and the percentage of footprints’ P-sites belonging to each bin in for each gene was calculated and plotted. Grey dashed line signifies a theoretical number where each bin of CDS contains an equivalent number of footprints for a sum of 100% across all 4 bins. **c**-**d**, Difference between footprint abundance in IP and that in Total ribosomes for each footprint size and bin (calculated in part a-b). **a** and **c,** Data from strains expressing WT+EV, *FLAG-UPF1*, *FLAG-UPF2,* or *FLAG-UPF3*. **b** and **d,** Data from strains expressing WT+EV, *UPF1-FLAG*, *UPF1-FLAG*/*upf2Δ*, or *upf1-DE572AA-FLAG*.

**Extended Data Figure 6.**
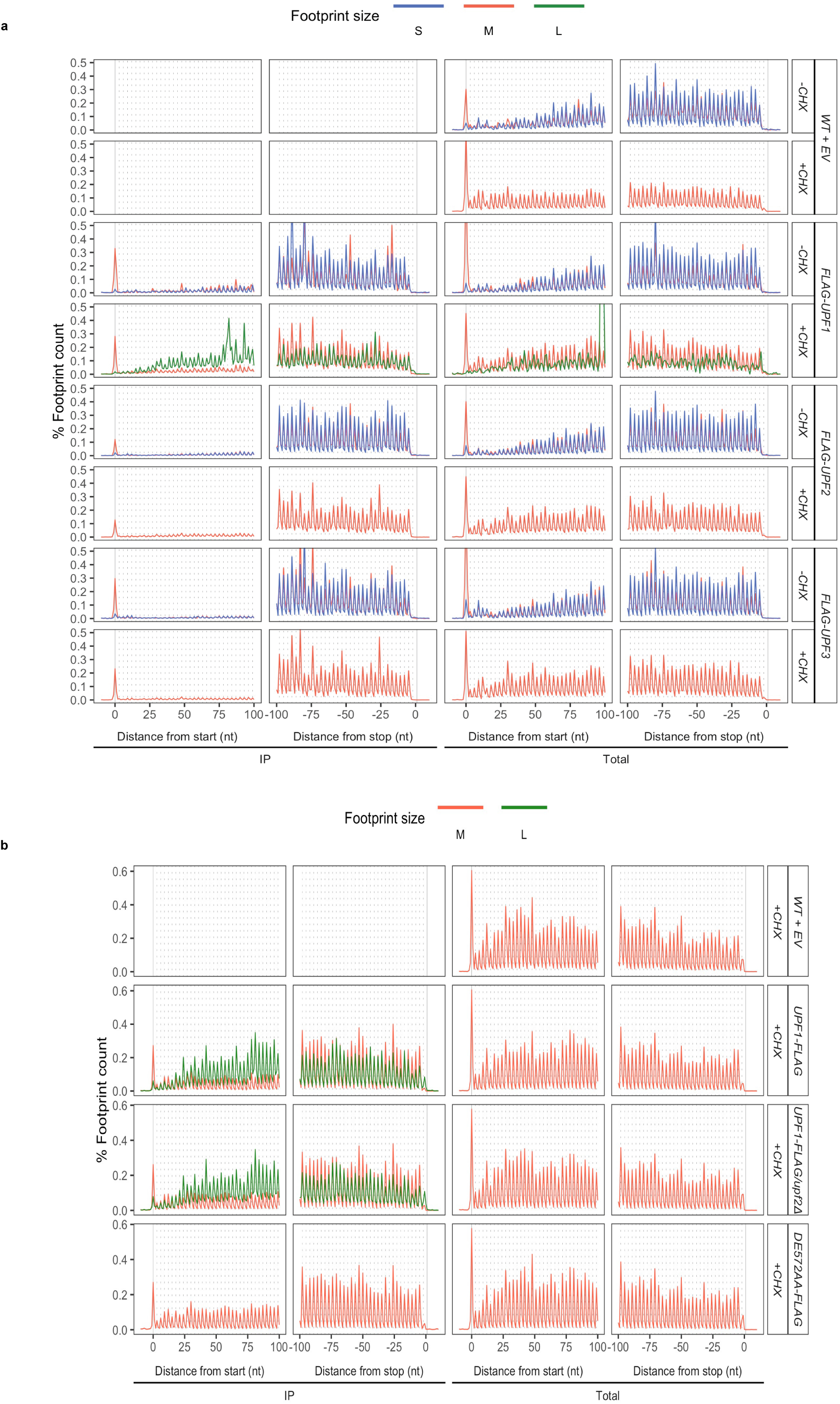
Abundance of ribosome footprints relative to canonical start or stop codons for immunopurified (IP) or total ribosomes (total) **a,** WT+EV, *FLAG-UPF1*, *FLAG-UPF2*, *UPF1-FLAG*/*upf2Δ*, or WT+EV strains treated with CHX**. b,** WT+EV, *UPF1-FLAG*, *UPF1-FLAG*/*upf2Δ*, or *upf1*-*DE572AA-FLAG* strains treated with CHX. For each footprint size, the number of footprint P-sites at each nucleotide position relative to start or stop codon of an mRNA is normalized to the total number of footprints of that size in the library and converted to percentage. The percentage of footprint counts in the first and last 100 nt of mRNAs were plotted. mRNAs whose coding region is less than 200 nt in length were disregarded from the plot to avoid repeated counting. Vertical grey solid lines signify the first nucleotide of the start and stop codons. Vertical grey dashed lines occur every 3 nucleotides to indicate reads in-frame with the start codon.

**Extended Data Figure 7.**
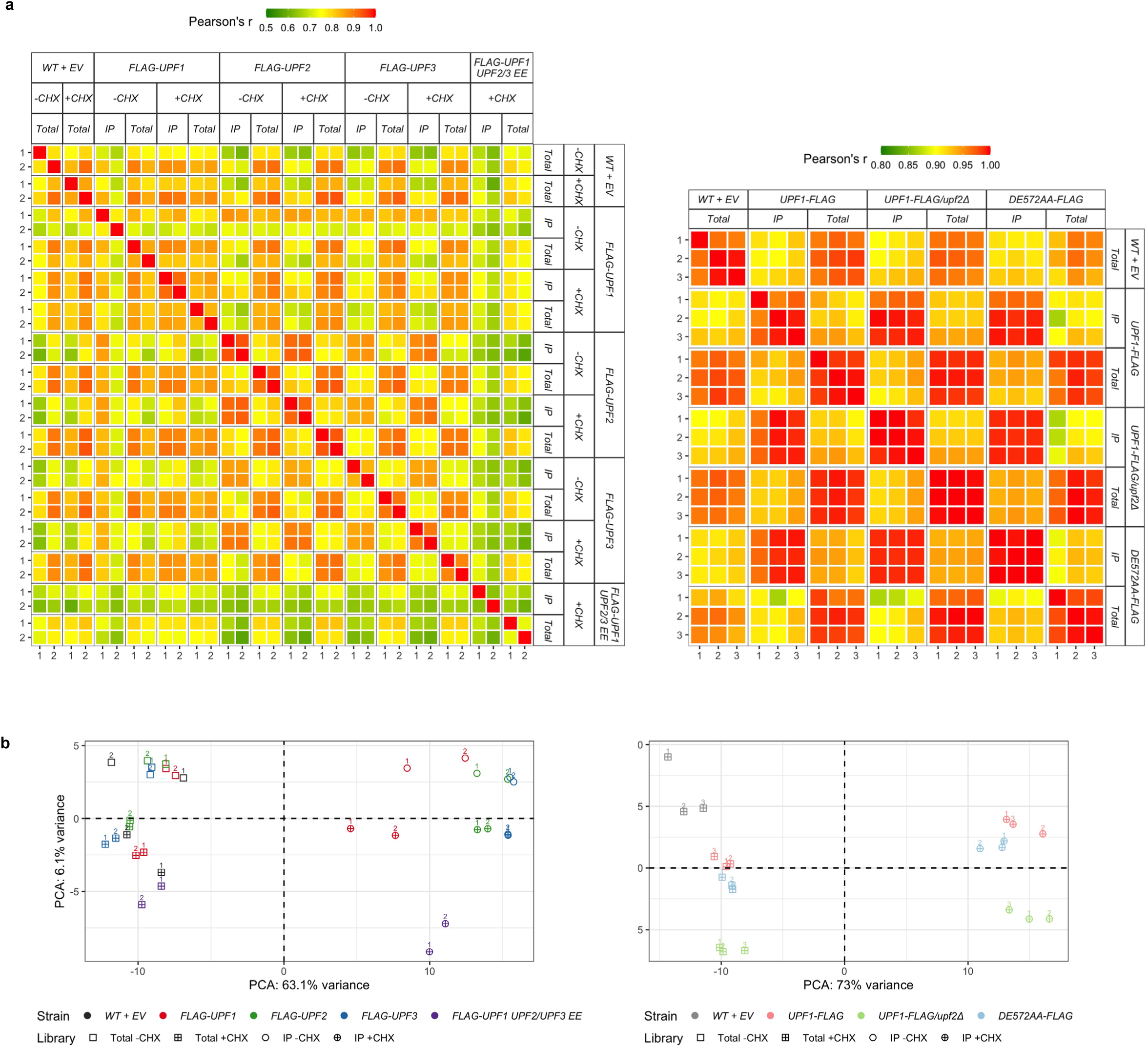
Reproducibility between replicates of ribosome profiling libraries. **a,** Correlation matrices showing pairwise Pearson’s correlation (*r*) of transcript abundance (log_10_-transformed, non-zero RPKM values) between total and IP ribosome profiling libraries from WT+EV, *FLAG-UPF1*, *FLAG-UPF2, FLAG-UPF3,* and *FLAG-UPF1 UPF2/UPF3 EE* (episomally expressed) strains (left) or WT+EV, *UPF1-FLAG*, *UPF1-FLAG*/*upf2Δ*, or *upf1*-*DE572AA-FLAG* strains (right). **b,** Principal Component Analyses (PCA) on the main footprint abundance (S and M for -CHX libraries; M for +CHX libraries) show abundance variation across multiple libraries. Libraries with smaller distance to one another are more similar than ones that are further apart. Variation in transcript abundance between samples are mostly explained by whether Upf factor-associated ribosomes were selectively recovered, as IP and Total libraries are segregated along the x-axis. Profiling libraries from WT+EV, *FLAG-UPF1*, *FLAG-UPF2, FLAG-UPF3,* and *FLAG-UPF1 UPF2/UPF3 EE* (episomally expressed) strains (left) or WT+EV, *UPF1-FLAG*, *UPF1-FLAG*/*upf2Δ*, or *upf1*-*DE572AA-FLAG* (right).

**Extended Data Figure 8.**
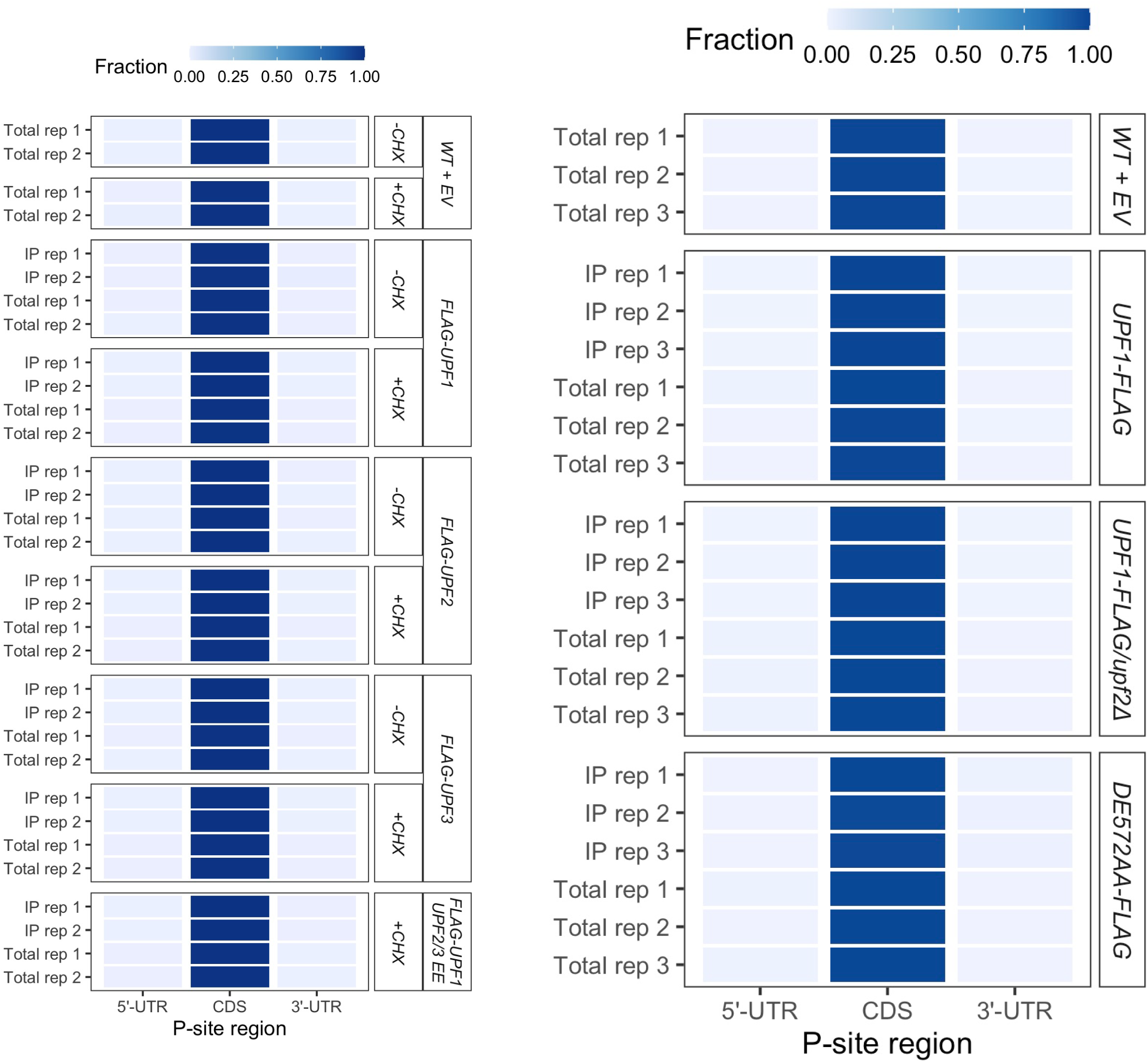
Fractions of mRNA regions (5’-UTR, coding, 3’-UTR) in which ribosome footprints’ P-site are located for each library. P-site location of each read was determined by testing whether its coordinate was located before the coordinate of the first nucleotide of the start codon (5’-UTR region), after the coordinate of the last nucleotide of the penultimate codon (3’-UTR region), or at or in between the two mentioned coordinates (coding region, CDS). The number of P-sites in each of the 3 regions was tabulated and normalized to the total number of reads in the library.

**Extended Data Figure 9.**
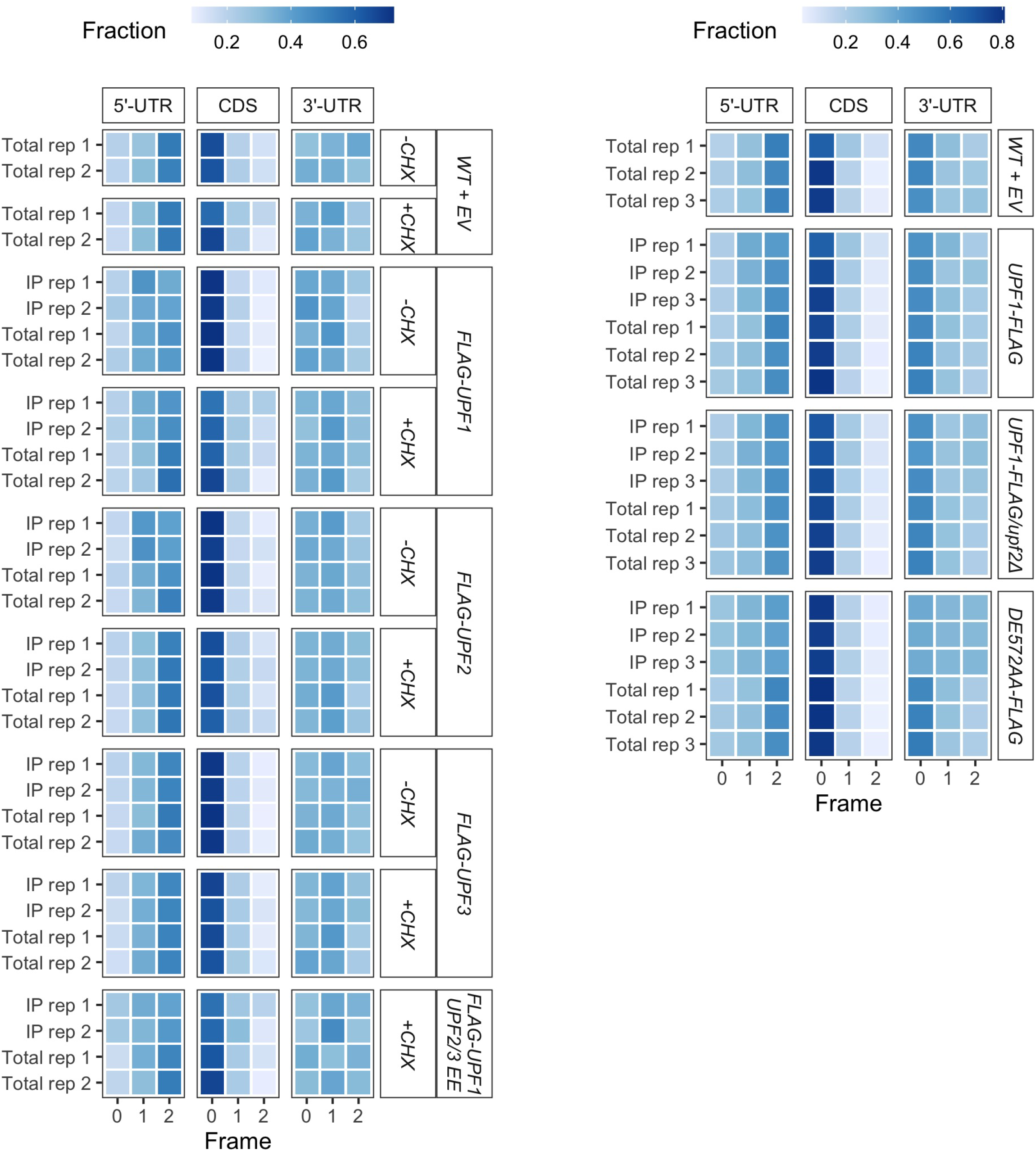
Fractions of ribosome footprints in each of the three reading frames in different mRNA regions (5’-UTR, coding, 3’-UTR). The reading frame of each read was calculated by dividing its coordinate relative to the first nucleotide of the start codon then recording the remainder. Thus, reading frame “0” refers to translation that is in-frame with the start codon. The number of reads in each reading frame in a particular mRNA region was tabulated and normalized to the total number of reads in that mRNA region.

**Extended Data Figure 10.**
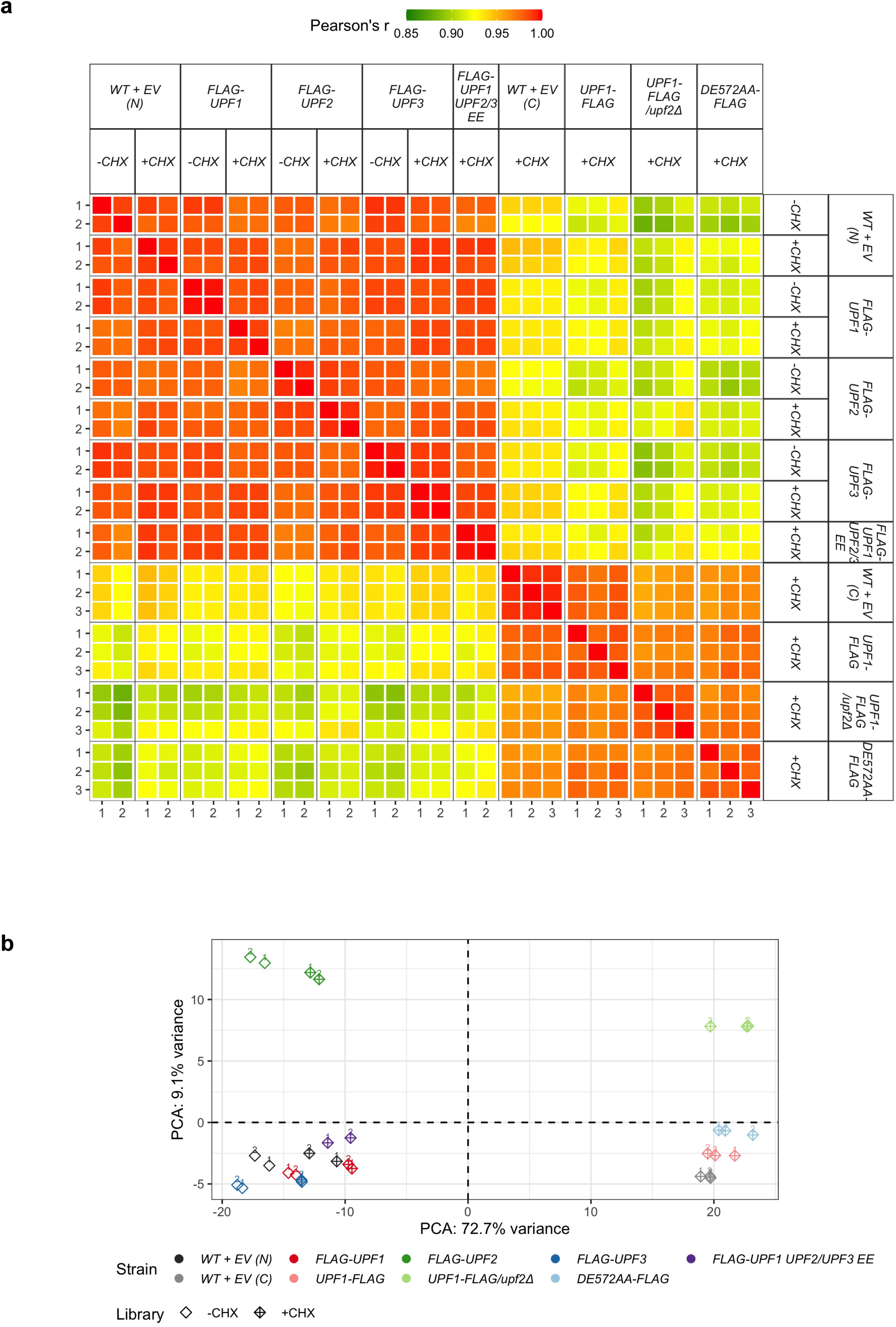
Reproducibility between replicates of RNAseq libraries. **a,** Correlation matrix showing pairwise Pearson’s correlation (*r*) of transcript abundance (log_10_-transformed, non-zero RPKM values) between RNAseq libraries. **b,** Principal Component Analysis (PCA) on reads abundance shows variation across multiple libraries. Libraries with smaller distance to one another are more similar than ones that are further apart. Variations in gene expression between strains are mostly explained by the difference in library preparation methods between N- and C-terminal FLAG-tagged strains (sample segregation along the x-axis) followed by the overexpression or deletion of *UPF2* (sample segregation along the y-axis).

## SUPPLEMENTARY INFORMATION

**Supplementary Table 1.** NMD substrate enrichment

**Supplementary Table 2.** L vs M abundance

**Supplementary Table 3.** Strains

**Supplementary Table 4.** Oligonucleotides

**Supplementary Table 5.** Plasmids

**Supplementary Table 6.** Sequencing statistics

**Supplementary Table 7.** P-site offsets

**Supplementary Table 8.** Mass spectrometry N-terminal data

**Supplementary Table 9.** Mass spectrometry C-terminal data

**Supplementary Table 10.** Mass spectrometry 6XHis-Upf1 data

**Supplementary Table 11.** Mass spectrometry FLAG-Upf1 Upf2-Upf3 EE data

